# Local administration of a novel siRNA modality into the CNS extends survival and improves motor function in the SOD1^G93A^ mouse model for ALS

**DOI:** 10.1101/2023.02.27.530262

**Authors:** Chunling Duan, Moorim Kang, Xiaojie Pan, Zubao Gan, Vera Huang, Guanlin Li, Robert F. Place, Long-Cheng Li

**Author notes:** Both authors contributed equally to this work (Chunling Duan, Moorim Kang).

## Abstract

Antisense oligonucleotides (ASOs) were the first modality to pioneer targeted gene knockdown in the treatment of ALS caused by mutant superoxide dismutase 1 (SOD1). RNA interference (RNAi) is another mechanism of gene silencing with historically superior potency in which short interfering RNAs (siRNAs) guide the RNA-induced silencing complex (RISC) to cleave complementary transcripts. However, delivery to extrahepatic tissues like the central nerve system (CNS) has been a bottleneck in the clinical development of RNAi. Herein, we identify potent siRNA duplexes for the knockdown of human SOD1 (hSOD1) in which medicinal chemistry and conjugation to an accessory oligonucleotide (ACO) enables durable and potent activity in CNS tissues. Local delivery via intracerebroventricular (ICV) or intrathecal (IT) injection into SOD1^G93A^ mice delayed disease progression and extended animal survival with superior efficacy compared to an ASO compound resembling Tofersen in sequence and chemistry. Treatment also prevented disease-related declines in motor function including improvements in animal mobility, muscle strength, and coordination. The ACO itself does not target any specific complementary nucleic acid sequence; rather, it imparts benefits conducive to bioavailability and delivery through its chemistry. The complete conjugate (*i.e*., siRNA-ACO) represents a novel modality for delivery of RNAi to the CNS in which we aim to pursue ALS as an exemplary indication for clinical development.

## INTRODUCTION

Amyotrophic lateral sclerosis (ALS) is a devastating neurodegenerative disease caused by the dysfunction and loss of motor neurons in the central nervous system (CNS). Disease progression is generally quick with a median survival at around 3-5 years following diagnosis (1). Early symptoms typically include muscle cramps, twitching, weakness, and stiffness. Patients inevitably begin to experience problems with movement and speech, which eventually manifests into assisted breathing, paralysis, and inevitable death.

Over 50 genes have been linked to ALS in which about 20% of all genetically defined cases are associated with mutation to the SOD1 gene (2). While SOD1 normally functions to protect cells from reactive oxygen species by catalyzing superoxide anions, loss of its activity has been shown to be independent of neuronal cell death (3, 4). Rather, mutant SOD1 confers a toxic gain-of-function caused by intracellular aggregation of misfolded protein that leads to motor neuron degradation (5). As such, reduction of SOD1 in CNS tissue has been the preferred therapeutic strategy for the treatment of ALS (6). For instance, Tofersen (BIIB067) is an antisense oligonucleotide (ASO) under clinical investigation that has shown therapeutic benefit in the treatment of ALS by suppressing mutant SOD1 mRNA levels (7).

RNA interference (RNAi) is another mechanism of gene silencing that has historically provided potent knockdown of targeted mRNA transcripts in comparison to other modalities (*i.e*., ASOs) (8). However, delivery to extrahepatic tissues (*i.e*., CNS) has limited its therapeutic development to broader indications. RNAi canonically requires double-stranded RNAs referred to as small interfering RNAs (siRNA) to guide the RNA-induced silencing complex (RISC) to complementary transcripts and suppress expression via mRNA cleavage (9). Disparate tolerances to medicinal chemistry advantageous for ASO delivery has been one of the reasons hindering drug development of RNAi for CNS indications (10–12). Only recently have new technologies emerged demonstrating siRNA delivery to CNS tissues with drug-like properties (13, 14).

We have developed another approach for delivering siRNA to the CNS by conjugating a single-stranded accessory oligonucleotide (ACO) to siRNA (siRNA-ACO) in a platform technology branded as SCAD (smart chemistry aided delivery). The ACO does not target any specific complementary nucleic acid sequence; rather, it imparts benefits conducive to bioavailability and delivery through its physiochemical composition (*i.e*., chemical modification and structure) typically incompatible with canonical siRNA duplexes. Herein, we identify potent siRNAs via high throughput screening (HTS) that knockdown human SOD1 in which medicinal chemistry and ACO conjugation enable durable activity for at least 8 weeks following local injection into the CNS of SOD1^G93A^ mice. Either intracerebroventricular (ICV) or intrathecal (IT) administration delays disease progression and extends animal survival in a dose-dependent manner. Furthermore, siRNA-ACO treatment prevented loss of motor function, as well as demonstrated superior efficacy in comparison to an ASO compound resembling Tofersen in both sequence and chemistry.

## RESULTS

### Development of siRNA drug candidates for knockdown of human SOD1

Sequence comprising the open reading frame (ORF) of human SOD1 transcript served as the template for siRNA design. A total of 268 siRNA duplexes were designed/synthesized at 21 nucleotides (nt) in length with no more than 4 repetitive nucleotides in a row and GC content between 35-65%. Knockdown activity of each siRNA was assessed in 293A cells using high throughput RT-qPCR at both 0.1 and 10 nM concentrations. Data was ranked according to mean knockdown activity in which 121 siRNAs reduced SOD1 by more than 90% at 10 nM (**Figure 1A)**. As an indicator of potency, 69 and 15 siRNAs reduced SOD1 levels at 0.1 nM treatments by over 50% and 75%, respectively. The overall top 30 performing siRNAs were subjected to an additional round of screening at 6 concentrations (*i.e*., 0.0064 - 20 nM) in 293A cells to demonstrate dose dependent activity in which propidium iodide (PI) was integrated into sample preparation to monitor variation in nucleic acid content as an indicator of untoward cytotoxicity (**Supplementary Figure 1**). Shown in **Figure 1B** is data for only the top 5 siRNAs (*i.e*., siSOD1-063, 047, 104, 005, and 258) with the most potent knockdown activity in absence of overt cytotoxicity (*i.e*., <20% reduction in PI staining).

**Figure 1.**
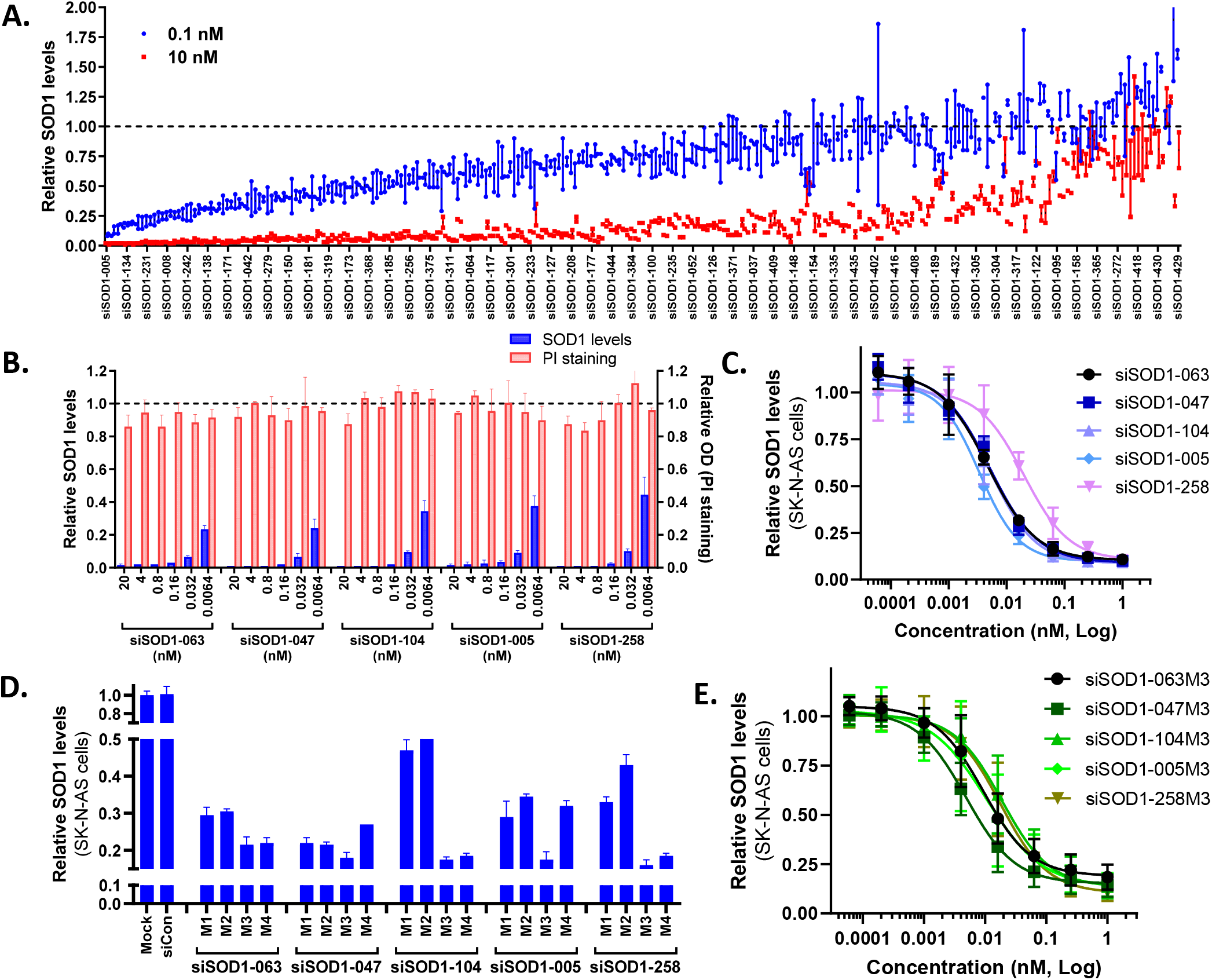
siRNA screen for SOD1 knockdown *in vitro*. **A**. 293A cells were transfected with each siRNA duplex (268 in total) in duplicate at 10 or 0.1 nM concentrations for 24 hours. SOD1 expression levels were quantified via RT-qPCR using gene specific primer sets. TBP was amplified as an internal reference used to normalize expression data. Shown are the expression values of SOD1 mRNA of each experimental replicate relative to Mock treatments (dotted line). Mock samples were transfected in absence oligonucleotide. **B**. Knockdown activity and cell viability of the top performing siRNAs was quantified in 293A cells at 6 escalating concentrations (*i.e*., 0.0064, 0.032, 0.16, 0.8, 4, and 20 nM) via RT-qPCR and PI staining, respectively. Shown are the results for both SOD1 knockdown and cytotoxicity of the top 5 performing siRNAs (*i.e*., siSOD1-063, 047, 104, 005, and 258) relative to Mock treatments (dotted line). **C**. Dose response curves were generated in SK-N-AS cells for each of the top 5 siRNAs at 8 treatment concentrations (*i.e*., 0.00006, 0.0002, 0.001, 0.004, 0.016, 0.063, 0.25, and 1 nM) via RT-qPCR. Data represents mean ± SD from 2 experimental replicates. **D**. SK-N-AS cells were transfected at 0.1 nM with 4 different chemically modified variants of each siRNA (*i.e*., M1, M2, M3, or M4) or non-specific siRNA control (siCon) for 24 hours. Knockdown activity was assessed via RT-qPCR relative to Mock treatment. **E**. Dose response curves were generated in SK-N-AS cells for each of the M3 modified variants (*i.e*., siSOD1-063M3, 047M3, 104M3, 005M3, and 258M3) via RT-qPCR. Data represents mean ± SD from 3 experimental replicates.

Dose response curves were subsequently generated for each of the top 5 siRNA candidates to validate potency in model cell lines representative of neuronal disease including SK-N-AS (**Figure 1C**) and T98G (**Supplementary Figure 2**) cells. As summarized in **Supplementary Table 1**, *in vitro* potency for each duplex was in the low picomolar range for both cell lines. Untoward cytotoxicity was also evaluated 72 hours after treatment at concentrations well above 200X the extrapolated IC_50_ values. As shown in **Supplementary Figure 3A-B**, only siSOD1-047 and 005 had no detectable impact on apoptosis or cell number in either SK-N-AS or T98G cells, whereas the remaining candidates (*i.e*., siSOD1-063, 104, and 258) had dose-dependent responses with regards to caspase 3/7 activity in T98G cells that inversely correlated with cell viability. While SK-N-AS cells appeared more tolerant to treatment, a similar pattern was observed for cell viability.

Several medicinal chemistry patterns referred to as M1, M2, M3, or M4 representing different duplex lengths (*i.e*., 20, 21, 22, and 23 nt, respectively) comprised of phosphorothionate (PS) backbone modifications at select positions with 2’-O-methylation (2’Ome) or 2’-fluoro (2’F) substitutions at every nucleotide were applied to each lead candidate and screened for target mRNA knockdown activity. As shown in **Figure 1D**, all M3 variants (*i.e*., siSOD1-063M3, 047M3, 104M3, 005M3, and 258M3) generally had better knockdown activity compared to the other chemically modified siRNAs at 0.1 nM treatment concentrations in SK-N-AS cells. A near identical pattern was also observed in T98G cells (**Supplementary Figure 4)**. To further characterize potency, dose response curves were generated for all M3-modified duplexes in both SK-N-AS (**Figure 1E**) and T98G (**Supplementary Figure 5**) cell lines. As summarized in **Supplementary Table 1**, *in vitro* potency was generally well-retained following chemical modification.

Accessory oligonucleotide (ACO) conjugation was developed to impart self-delivery properties similar to ASOs by sharing medicinal chemistry normally not tolerated by canonical siRNAs. As such, the passenger strand of each M3 variant (*i.e*., siSOD1-063M3, 047M3, 104M3, 005M3, and 258M3) was synthesized covalently linked to a 14-nucleotide ACO (referred to AC1) via a short PEG linker (L9) that possessed PS backbone substitutions and 2’-O-methoxyethyl (2’MOE) modifications at every position within the ACO (**Figure 2A**). Gel shift assays demonstrate that AC1 conjugation promotes novel interactions with factors in human serum compared to only chemically modified duplexes (**Supporting Figure 6**). AC1 was intentionally designed to lack significant homology to any known transcript with chemistry typically incompatible with ASO-mediated RNase H activity. As shown in **Figure 2B**, AC1 treatment alone had no detectable knockdown of SOD1 at 0.25 and 2.5 nM concentrations *in vitro*, whereas activity was only perceived when conjugated to siRNA. Additional dose response analysis noted a ~10X loss in siRNA potency as a consequence of AC1 conjugation (*i.e*., siSOD1-005M3-AC1) compared to only chemically modified duplex (*i.e*., siSOD1-005M3) (**Figure 2C**). However, potency was restored upon modification with 5’-(*E*)-vinylphosphonate (5’VP) at the 5’ terminus of the guide strand (*i.e*., siSOD1-005M3-AC1^VP^) (**Figure 2D**). In comparison to an ASO resembling Tofersen in both sequence and chemistry (*i.e*., ASO^SOD1^), SOD1 knockdown with either siRNA-ACO variant (*i.e*., siSOD1-005M3-AC1 or siSOD1-005M3-AC1^VP^) was still more potent than ASO^SOD1^ regardless of 5’VP modification (**Figure 2D**).

**Figure 2.**
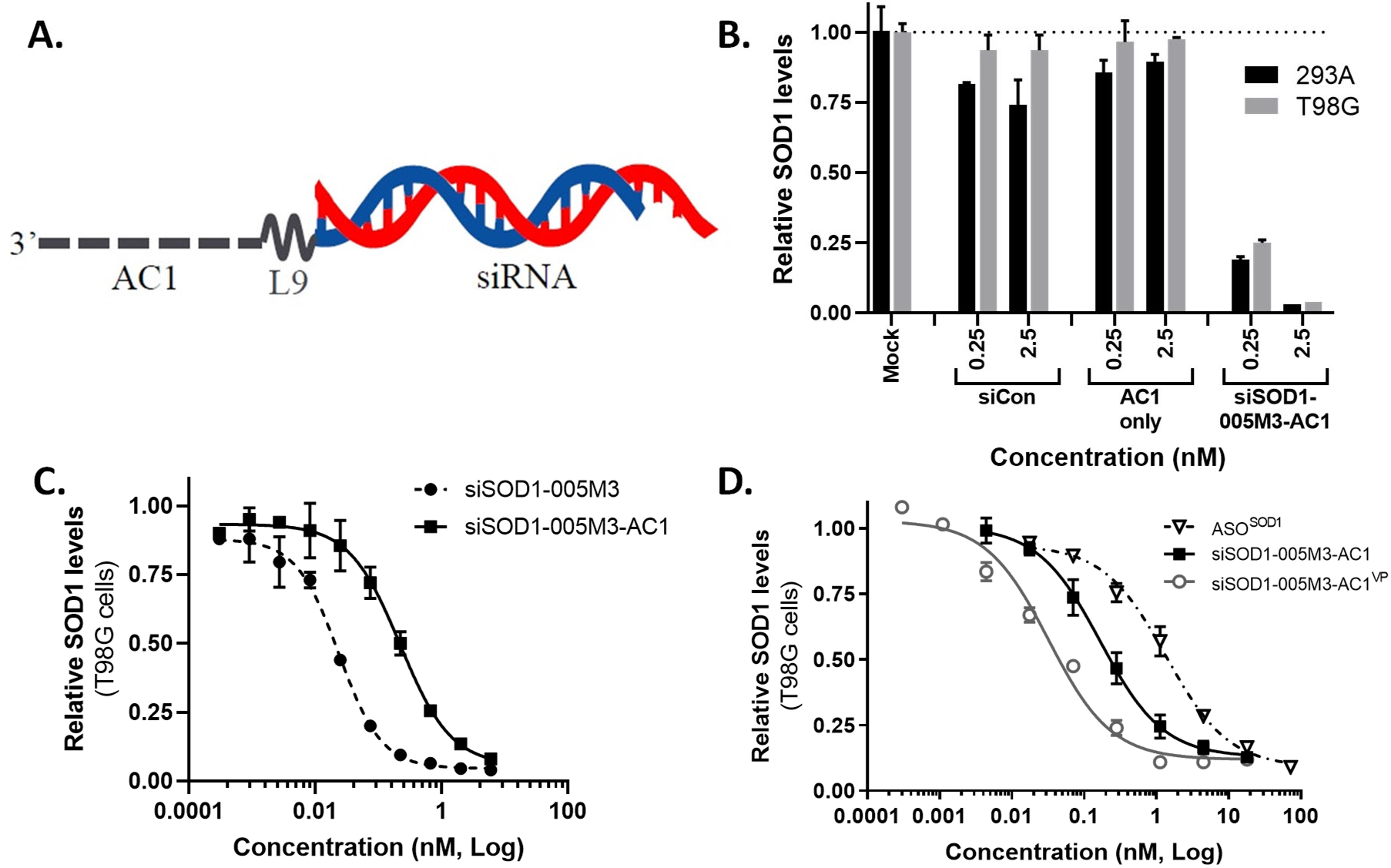
siRNA-ACO knockdown activity *in vitro*. **A**. Depicted is a visual representation of the siRNA-ACO structure in which a 14-nt ACO (referred to AC1) is conjugated to the 3’-terminus of the passenger strand via short PEG linker (L9). **B**. 293A and T98G cells were transfected at 0.25 or 2.5 nM concentrations with exemplary siRNA-ACO (*i.e*., siSOD1-005M3-AC1) or AC1 only for 24 hours. Mock samples were transfected in absence oligonucleotide. Treatment with siCon served as a negative control for knockdown activity. SOD1 expression levels were quantified via RT-qPCR using gene specific primer sets. TBP was amplified as an internal reference used to normalize expression data. Shown are the expression values of SOD1 mRNA of each experimental replicate relative to Mock treatments (dotted line). **C**. Dose response curves were generated in T98G cells comparing siRNA knockdown activity with (siSOD1-005M3-AC1) or without (siSOD1-005M3) AC1 conjugation at 10 treatment concentrations (*i.e*., 0.0003, 0.0009, 0.0027, 0.0082, 0.024, 0.074, 0.22, 0.67, 2, and 6 nM) via RT-qPCR. **D**. Dose response curves were generated comparing knockdown activity of siRNA-ACOs with (siSOD1-005M3-AC1^VP^) or without (siSOD1-005M3-AC1) 5’VP modification to ASO^SOD1^.

siRNA-ACO conjugates were subsequently synthesized using M3 chemistry and 5’VP modification (*i.e*., siSOD1-063M3-AC1^VP^, 047M3-AC1^VP^, 104M3-AC1^VP^, 005M3-AC1^VP^, and 258M3-AC1^VP^) for downstream screening *in vivo*. Prior to treatment, dose response curves were generated in both SK-N-AS and T98G cells to validate activity of each siRNA-ACO candidate (**Supplementary Figure 7A-B**). As summarized in **Supplementary Table 1**, *in vitro* potencies were generally well conserved in comparison to their non-conjugate forms. Apoptosis and cell viability were also quantified 72 hours after treatments to measure any changes in cytotoxicity as a consequence of chemical modification and ACO conjugation. As shown in **Supplementary Figure 8A-B**, siSOD1-047M3-AC1^VP^ and 005M3-AC1^VP^ remained generally unaffected with nominal impact on cell health in both SK-N-AS and T98G cells, whereas siSOD1-104M3-AC1^VP^ retained signs of untoward cytotoxicity and the remaining candidates (*i.e*., siSOD1-063M3-AC1^VP^ and 258M3-AC1^VP^) noted an improvement in *in vitro* safety (*i.e*., reduction in caspase 3/7 activity) compared to their non-modified forms (**Supplementary Figure 3A-B**).

### *In vivo* selection of siRNA-ACO drug candidates

Mice hemizygous for SOD1^G93A^ transgene express a mutant form of human SOD1 and exhibit disease phenotypes similar to ALS including progressive loss of motor function and abbreviated life span via neuronal degradation (15, 16). To test knockdown activity *in vivo*, SOD1^G93A^ mice were treated via ICV injection at 10 nmole/dose of each siRNA-ACO conjugate (*i.e*., siSOD1-063M3-AC1^VP^, 047M3-AC1^VP^, 104M3-AC1^VP^, 005M3-AC1^VP^, and 258M3-AC1^VP^). All siRNA-ACOs were formulated in aCSF in which treatment alone served as a vehicle control to establish baseline expression, while siCON1-AC1^VP^ functioned as a negative control for siRNA-ACO activity. Mice were also treated with ASO^SOD1^ at ~3X molar excess of siRNA-ACO (*i.e*., 28 nmole/dose) as a comparative control for hSOD1 knockdown. Tissues from the CNS (*i.e*., frontal cortex, cerebellum, cerebrum, and spinal cord) and periphery (*i.e*., liver) were harvested on day 14 after treatment in which hSOD1 expression levels were quantified via RT-qPCR. As shown in **Figure 3A**, both siSOD1-047M3-AC1^VP^ and siSOD1-005M3-AC1^VP^ had more potent knockdown activity in the CNS tissues compared to the other siRNA-ACO candidates. Activity was well localized within the CNS in which drainage to peripheral tissue (*i.e*., liver) did not produce comparable activity. Knockdown via ASO^SOD1^ produced results similar to siSOD1-258M3-AC1^VP^, yet substantially less than the lead candidates (*i.e*., siSOD1-047M3-AC1^VP^ and 005M3-AC1^VP^).

**Figure 3.**
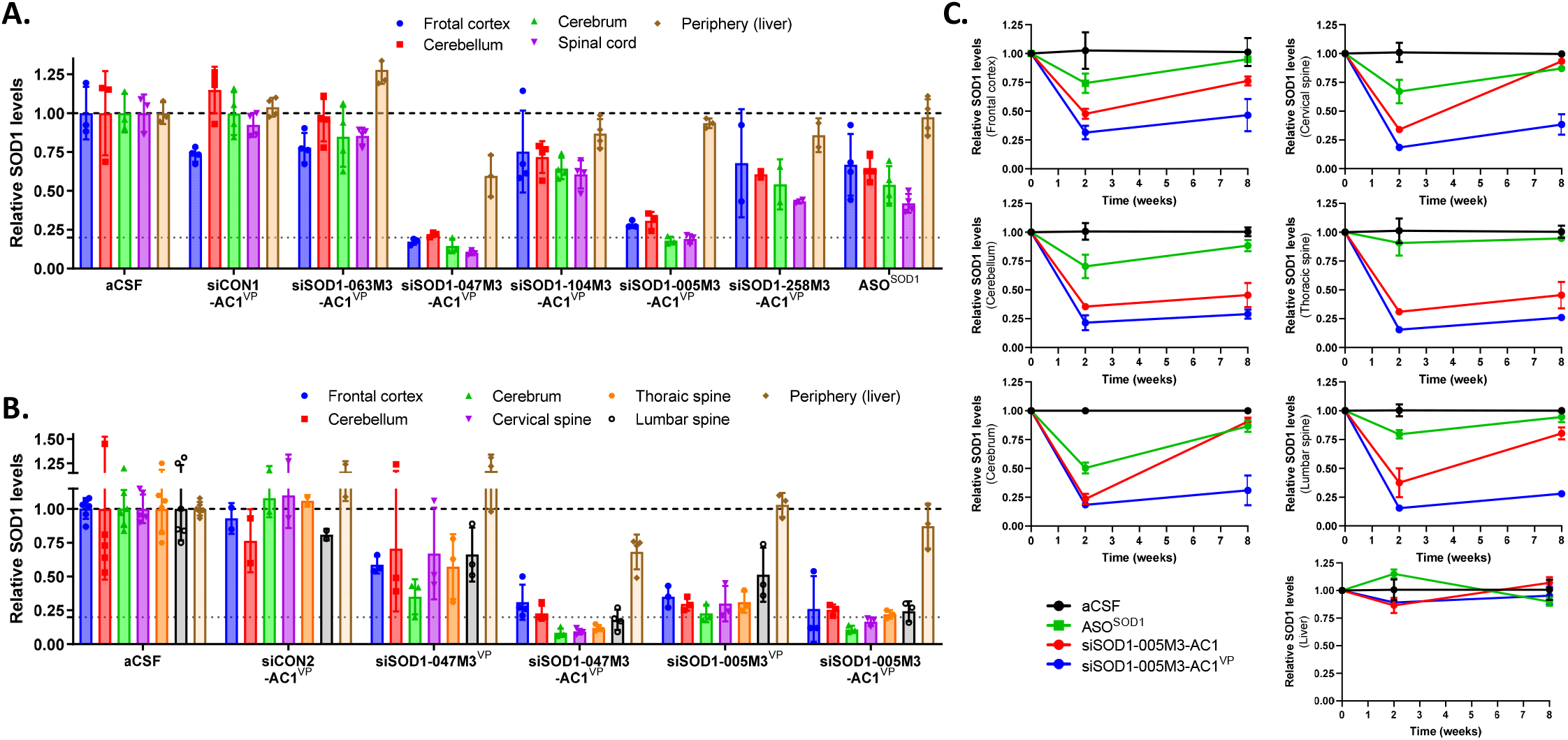
siRNA-ACO activity in CNS tissue of SOD1^G93A^ mice. **A**. Adult SOD1^G93A^ mice were treated via ICV injection with each siRNA-ACO drug candidate (*i.e*., siSOD1-063M3-AC1^VP^, 047M3-AC1^VP^, 104M3-AC1^VP^, 005M3-AC1^VP^, and 258M3-AC1^VP^) at 10 nmole. Animals administered ASO^SOD1^ were dosed at 28 nmole. Treatment with aCSF alone was used as a vehicle control to establish baseline expression, while a non-specific siRNA-ACO (*i.e*., siCON1-AC1^VP^) served as a negative control for knockdown activity. Human SOD1 expression was quantified via RT-qPCR using gene specific primer sets in tissues from the CNS (*i.e*., frontal cortex, cerebellum, cerebrum, and spinal cord) and periphery (*i.e*., liver) on day 14 after treatment. Mouse Tbp (mTbp) was amplified as an internal reference to normalize expression data. Shown are the mean expression values ± SD (n=3-4 mice/group) of hSOD1 relative to aCSF treatment. The dotted gray line represents 80% knockdown relative to baseline (dashed line). **B**. Knockdown activity was quantified in tissues from the brain (*i.e*., frontal cortex, cerebellum, and cerebrum), spinal cord (*i.e*., cervical, thoracic, and lumbar), and periphery (*i.e*.,liver) via RT-qPCR on day 14 following treatment via ICV injection at a fixed molecular dose (20 nmole) with siRNA-ACO candidates (*i.e*., siSOD1-047M3-AC1^VP^ and 005M3-AC1^VP^) or their non-conjugated derivatives (*i.e*., siSOD1-047M3^VP^ and 005M3^VP^). Treatment with siCON2-AC1^VP^ served as a negative control for knockdown activity. Shown are the mean expression values ± SD (n=2-6 mice/group) of hSOD1 relative to aCSF treatment. **C**. Knockdown durability was assessed via RT-qPCR in the indicated CNS tissues at 2 and 8 weeks following single dose of siSOD1-005M3-AC1, siSOD1-005M3-AC1^VP^, or ASO^SOD1^ at 20 mg/kg via ICV injection. Treatment with aCSF alone served as a vehicle control. Data represents mean ± SD (n=3-4 mice/group) relative to SOD1 levels pre-treatment (*i.e*., 0 weeks).

SOD1^G93A^ mice were also treated via ICV injection at equimolar quantities (*i.e*., 10 nmole/dose) with siRNA-ACOs (*i.e*., siSOD1-047M3-AC1^VP^ or siSOD1-005M3-AC1^VP^) in comparison to non-conjugate controls (*i.e*., siSOD1-047M3^VP^ or siSOD1-005M3^VP^) to demonstrate the effect AC1 conjugation imparts on knockdown activity *in vivo*. As shown in **Figure 3B**, both siRNA-ACO duplexes provided greater knockdown activity comparative to their non-conjugate cognates in all tissues of the brain (*i.e*., frontal cortex, cerebellum, and cerebrum) and spinal cord (*i.e*., cervical, thoracic, and lumbar spine) with minimal activity in the periphery (*i.e*., liver). Knockdown durability was also characterized for siSOD1-005M3-AC1^VP^ relative to its non-5’VP control (*i.e*., siSOD1-005M3-AC1) with comparison to ASO^SOD1^. As shown in **Figure 3C**, knockdown via siSOD1-005M3-AC1^VP^ was generally well sustained out to 8 weeks in all CNS tissues, whereas activity of siSOD1-005M3-AC1 and ASO^SOD1^ began to return to baseline. Taken together, the combination of both AC1 conjugation and 5’VP modification provided our siRNAs with enhanced activity needed for a durable response *in vivo*.

### Single dose ICV injection of siRNA-ACO delays disease progression and prolongs survival in SOD1^G93A^ mice

Adult SOD1^G93A^ mice were treated via ICV injection with lead siRNA-ACO candidates (*i.e*., siSOD1-047M3-AC1^VP^ or siSOD1-005M3-AC1^VP^) at 50, 100, 200, or 400 μg/dose on PND85 or 60, respectively. Drug concentrations and hSOD1 expression levels were quantified in a subset of animals on day 14 after treatment in CNS tissue (*i.e*., cerebellum, cerebrum, and spinal cord). As shown in **Figure 4A-B**, both activity and tissue accumulation of siSOD1-047M3-AC1^VP^ and siSOD1-005M3-AC1^VP^ were dose-dependent in which SOD1 knockdown inversely correlated with increasing concentrations of siRNA-ACO within the CNS tissues. Drug concentrations projected to elicit an ED_50_ (median effective dose) response in each tissue are summarized in **Supplementary Table 2**.

**Figure 4.**
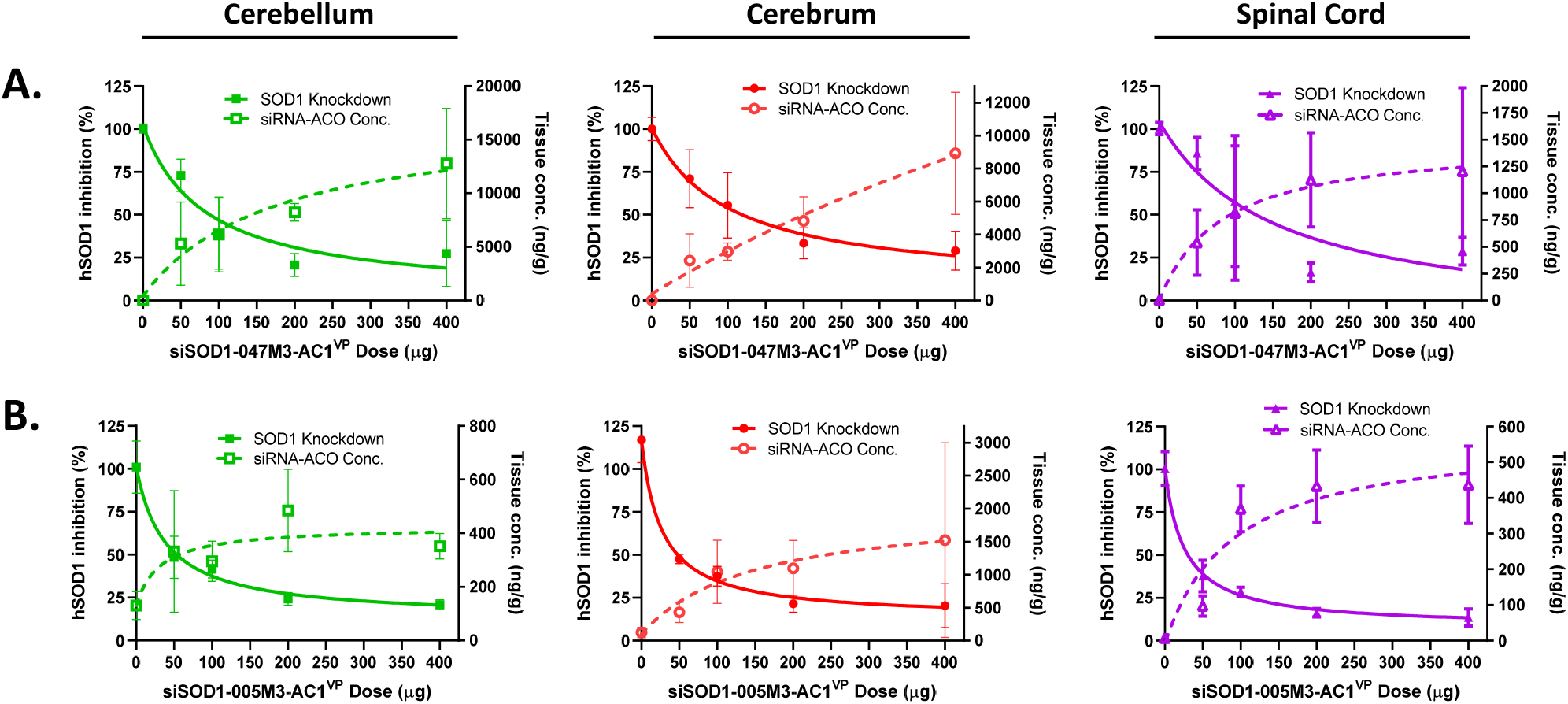
Dose-dependent relationship between knockdown activity and tissue accumulation of siRNA-ACO. Adult hSOD1^G93A^ mice were treated with siSOD1-047M3-AC1^VP^(**A**) or siSOD1-005M3-AC1^VP^ (**B**) at the indicated doses (*i.e*., 50, 100, 200, or 400 μg) via ICV injection. Knockdown activity of hSOD1 was quantified via RT-qPCR in select CNS tissues (*i.e*., cerebellum, cerebrum, and spinal cord) on day 14 following. Animals treated with aCSF alone represent expression levels at baseline and drug quantities detectable in absence of siRNA-ACO treatment (0 μg). Knockdown activity is shown as percent (%) inhibition of SOD1 relative to baseline (0 μg). Drug concentrations are shown as siRNA-ACO quantities relative to tissue sample mass (μg/g). Data represents mean ± SD (n=3-4 mice/group).

In remaining animals, changes in body weight were plotted to monitor growth rate and disease progression. As shown in **Figure 5A**, all groups treated with either siSOD1-047M3-AC1^VP^ or siSOD1-005M3-AC1^VP^ continued to gain weight in comparison to aCSF treatment. In addition, disease-related weight loss was delayed in a dose-dependent manner as noted by time needed for growth rates to return to starting weight (dotted line). Disease progression was confirmed in each animal when a 10% loss in peak body weight was recorded. Plotting data via Kaplan-Meier curves indicated when animals in each treatment group transitioned to progressive disease (**Figure 5B**). Data was collected until animals inevitably succumbed to their disease in which survival curves were also generated (**Figure 5C**). To summarize, both siSOD1-047M3-AC1^VP^ (**Table 1**) and siSOD1-005M3-AC1^VP^ (**Table 2**) treatment delayed disease progression and extended animal survival in which the highest dose (*i.e*., 400 μg) prolonged life by 70 and 111.5 days compared to vehicle controls, respectively.

**Figure 5.**
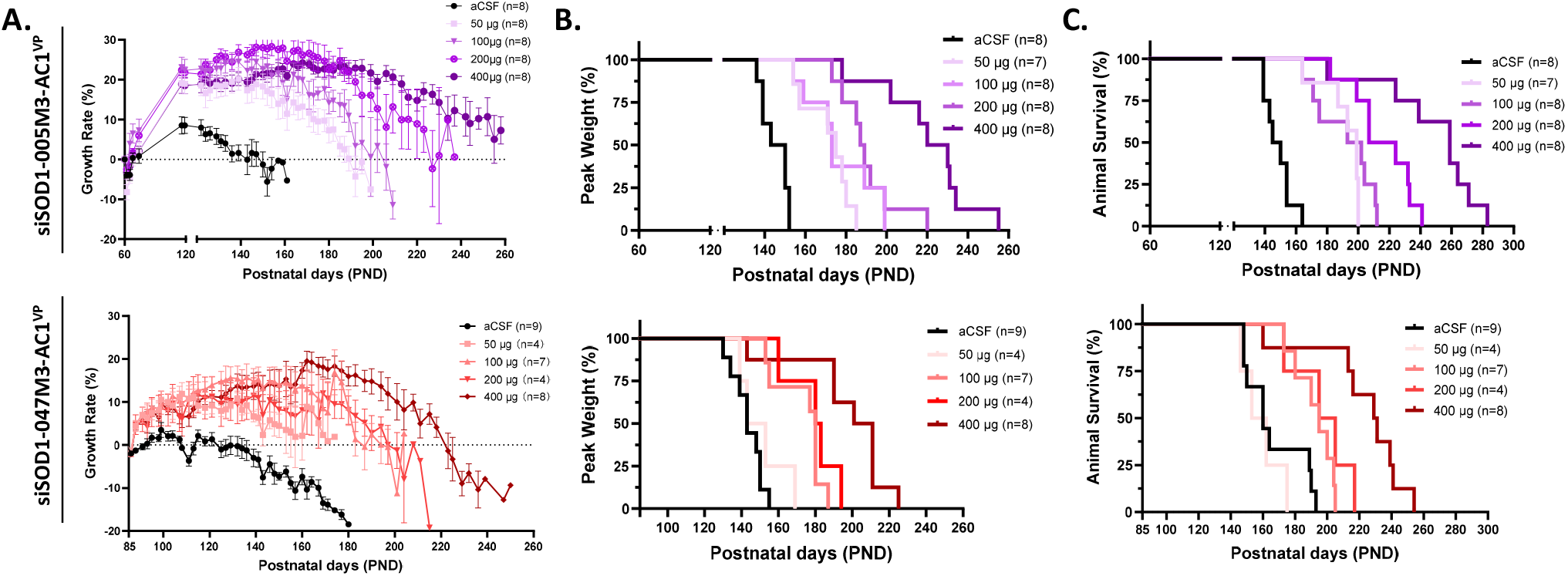
Single dose treatment via ICV injection with siRNA-ACO delays disease progression and prolongs survival. **A**. Adult hSOD1^G93A^ mice were treated with siSOD1-047M3-AC1^VP^ or siSOD1-005M3-AC1^VP^ at the indicated doses (*i.e*., 50, 100, 200, or 400 μg) via ICV injection on PND85 or PND60, respectively. Growth rates (*i.e*., percent change in body weight) relative to first day of treatment (dotted line) were plotted to monitor disease progression. **B**. Data is plotted as percent animals in each treatment group at peak body weight. **C**. Animal survival is plotted as percentage of surviving animals in each treatment group. Animal numbers (n) are indicated in each graph.

**Table 1.** Median age for disease onset and animal survival following single dose siSOD1-047M3-AC1^VP^ on PND85

| Treatment | aCSF | siSOD1-047M3-AC1<sup>VP</sup> |  |  |  |
| --- | --- | --- | --- | --- | --- |
| Dose (μg) | - | 50 | 100 | 200 | 400 |
| Animals No. (n) | 9 | 4 | 7 | 4 | 8 |
| **Disease Onset** |  |  |  |  |  |
| Median Age (Days) | 143 | 148 | 180* | 181.5† | 206* |
| **Survival** |  |  |  |  |  |
| Median Age (Days) | 160 | 157.5 | 195† | 200 | 230* |

\**P* < 0.001 compared to aCSF group
†*P* < 0.01 compared to aCSF group

**Table 2.** Median age for disease onset and animal survival following single dose siSOD1-005M3-AC1^VP^ on PND60

| Treatment | aCSF | siSOD1-005M3-AC1<sup>VP</sup> |  |  |  |
| --- | --- | --- | --- | --- | --- |
| Dose (µg) | - | 50 | 100 | 200 | 400 |
| Animals No. (n) | 8 | 7 | 8 | 8 | 8 |
| **Disease Onset** |  |  |  |  |  |
| Median Age (Days) | 146.5 | 175* | 173* | 188* | 225* |
| **Survival** |  |  |  |  |  |
| Median Age (Days) | 147.5 | 199* | 197.5* | 215.5* | 259* |

\* $P < 0.001$ compared to aCSF group

### Pathogenic mutation in target sites of siRNA-ACO impact lead selection

*In silico* analysis of pathogenic single nucleotide polymorphisms (SNPs) located within the target site of siSOD1-005M3-AC1^VP^ revealed 5 total SNPs in which 4 were in the region complementary to its “seed” sequence (**Supplementary Figure 9A**). Mismatches in this region are known to inhibit siRNA activity, which could eliminate a projected ~8.9-12.4% of the global ALS^SOD1^ patients from its treatment pool (17, 18). Conversely, siSOD1-047M3-AC1^VP^ has only 2 reported pathogenic SNPs within its target site (*i.e*., P.E22G and P.F21C) comprising approximately 4.04% and 2.70% of the global ALS^SOD1^ population, respectively (**Supplementary Figure 9B**). Luciferase reporter constructs (*i.e*., pLuc^SOD1^, pLuc^P.E22G^, and pLuc^P.F21C^) containing either consensus sequence perfectly complementary to siSOD1-047M3-AC1^VP^ guide strand or one of pathogenic mutations (*i.e*., P.E22G and P.F21C) were co-transfected into 239A cells along with siRNA-ACO. Knockdown of luciferase activity was specific to siSOD1-047M3-AC1^VP^ as transfection with a scramble control (*i.e*., siCON2-AC1^VP^) did not reduce reporter expression (**Supplementary Figure 9C**). Dose response data indicated that the P.E22G mutation does not have any significant impact on siSOD1-047M3-AC1^VP^ knockdown activity/potency compared to consensus target site sequence, while P.F21C was partially resilient to treatment demonstrating incomplete knockdown and inferior potency (**Supplementary Figure 9D)**.

### IT injection of siRNA-ACO provides therapeutic efficacy against ALS in SOD1^G93A^ mice

Based on perceived patient populations, siSOD1-047M3-AC1^VP^ was selected for further *in vivo* analysis. Male and female SOD1^G93A^ mice were treated via IT injection with two sequential doses of siSOD1-047M3-AC1^VP^ on PND 68 and 100 at 75, 150, or 300 μg/dose. A non-specific siRNA-ACO (*i.e*., siCON3-AC1^VP^) served as a negative control for therapeutic efficacy. Animal weight was monitored with comparison to wild-type animals (*i.e*., WT) in which all siSOD1-047M3-AC1^VP^ treatments provided similar benefit to male mice, whereas weight gain in females appeared more noticeably dose-dependent (**Figure 6A**). Both disease progression (*i.e*., 10% loss in peak body weight) and animal survival were also plotted in which siSOD1-047M3-AC1^VP^ treatments delayed advance disease and extended survival in both male and female mice (**Figure 6B-C**). **Table 3** summarizes median days for disease progression and survival following siRNA-ACO treatment via IT injection in male and female populations. Overall, siCON3-AC1^VP^ provided no therapeutic benefit in comparison to vehicle control, while equivalent doses of siSOD1-047M3-AC1^VP^ (*i.e*., 150 μg) extended animal survival in males and females by 61 and 29 days, respectively.

**Figure 6.**
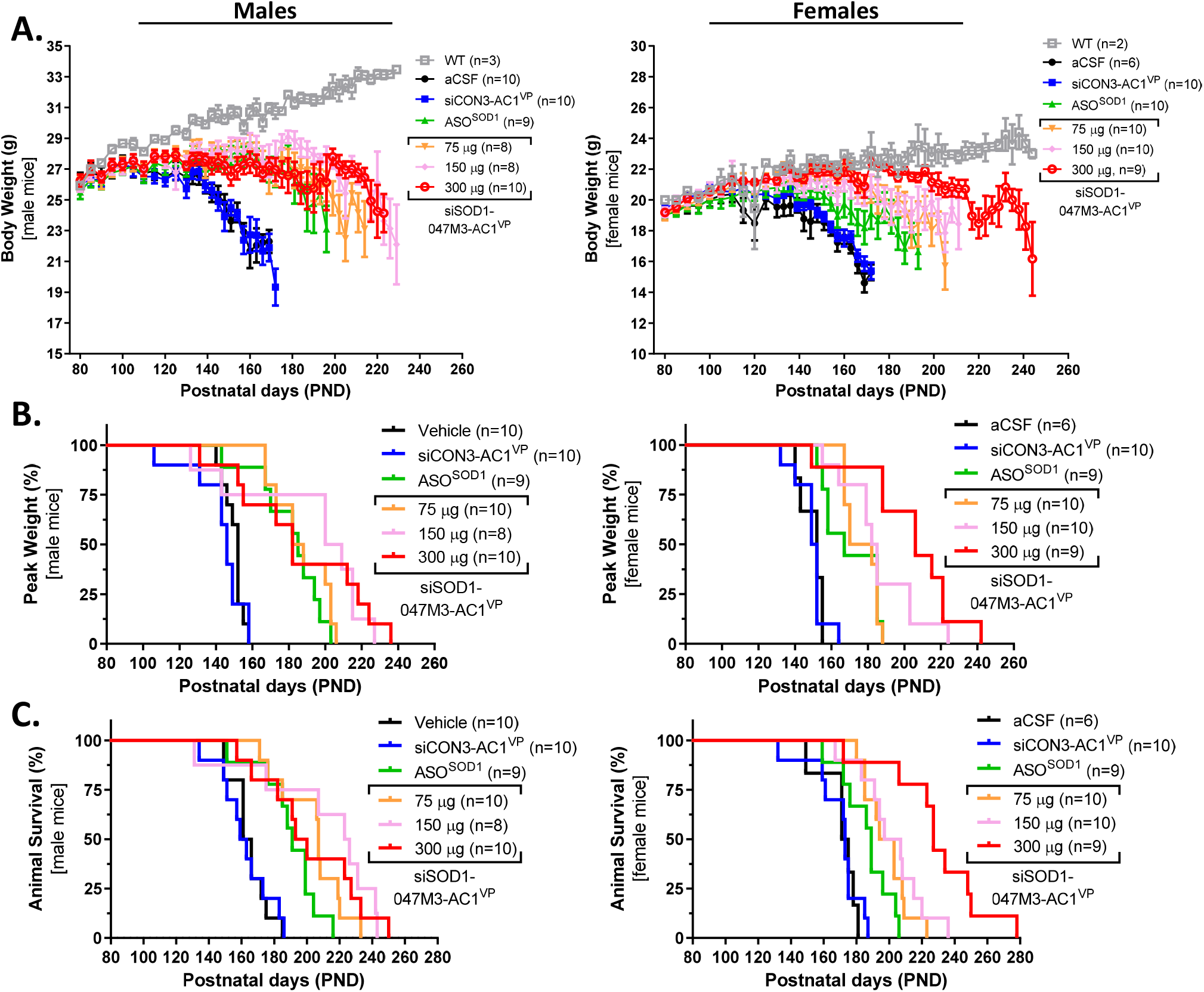
siRNA-ACO delays disease progression and prolongs survival via IT injection. **A**. Male and female adult hSOD1^G93A^ mice were treated twice with siSOD1-047M3-AC1^VP^ at the indicated doses (*i.e*., 75, 150, or 300 μg) via IT injection on PND68 and PND100. Non-specific control (*i.e*., siCON3-AC1^VP^) and ASO^SOD1^ were dosed at 150 μg/injection. Treatment with aCSF served as a vehicle control. Body weight in grams (g) was plotted comparative to background animals (WT) to monitor disease progression. **B**. Data is plotted as percent animals in each treatment group at peak body weight. **C**. Animal survival is plotted as percentage of surviving animals in each treatment group. Animal numbers (n) are indicated in each graph.

**Table 3.** Median age of onset and survival in male and female mice following siSOD1-047M3-AC1^VP^ treatment via IT injection

| Treatment | aCSF | siCON3-AC1<sup>VP</sup> | ASO<sup>SOD1</sup> | siSOD1-047M3-AC1<sup>VP</sup> |  |  |
| --- | --- | --- | --- | --- | --- | --- |
| Dose (μg) | - | 150 | 150 | 75 | 150 | 300 |
| **Animal Number (n)** |  |  |  |  |  |  |
| Male (♂) | 10 | 10 | 9 | 10 | 8 | 10 |
| Female (♀) | 6 | 10 | 9 | 10 | 10 | 9 |
| **Disease Onset (days)** |  |  |  |  |  |  |
| Male (♂) | 152 | 146 | 185* | 185* | 204.5† | 182† |
| Female (♀) | 152 | 150.5 | 167† | 176* | 183.5* | 206* |
| **Survival (days)** |  |  |  |  |  |  |
| Male (♂) | 163.5 | 161 | 191* | 207* | 224.5† | 196.5* |
| Female (♀) | 173 | 173 | 189 | 198.5* | 202* | 227* |

\**P* < 0.001 compared to aCSF group
†*P* < 0.01 compared to aCSF group

Neuromuscular performance was also evaluated in groups comprised of both male and female mice. Distance traveled via open field roaming (**Figure 7A**), rotarod test, (**Figure 7B**), and grip strength (**Figure 7C**) all showed siSOD1-047M3-AC1^VP^ treatment greatly improved motor function in SOD1^G93A^ mice, which was generally well sustained until the end of study. Comparing rotarod performance of each individual animal at an early timepoint prior to measurable weight loss (*i.e*., PND90) to their respective last timepoints further indicates siSOD1-047M3-AC1^VP^ treatment retained or improved neuromuscular performance in a majority of animals compared to controls or ASO^SOD1^ (**Supplementary Figure 10**).

**Figure 7.**
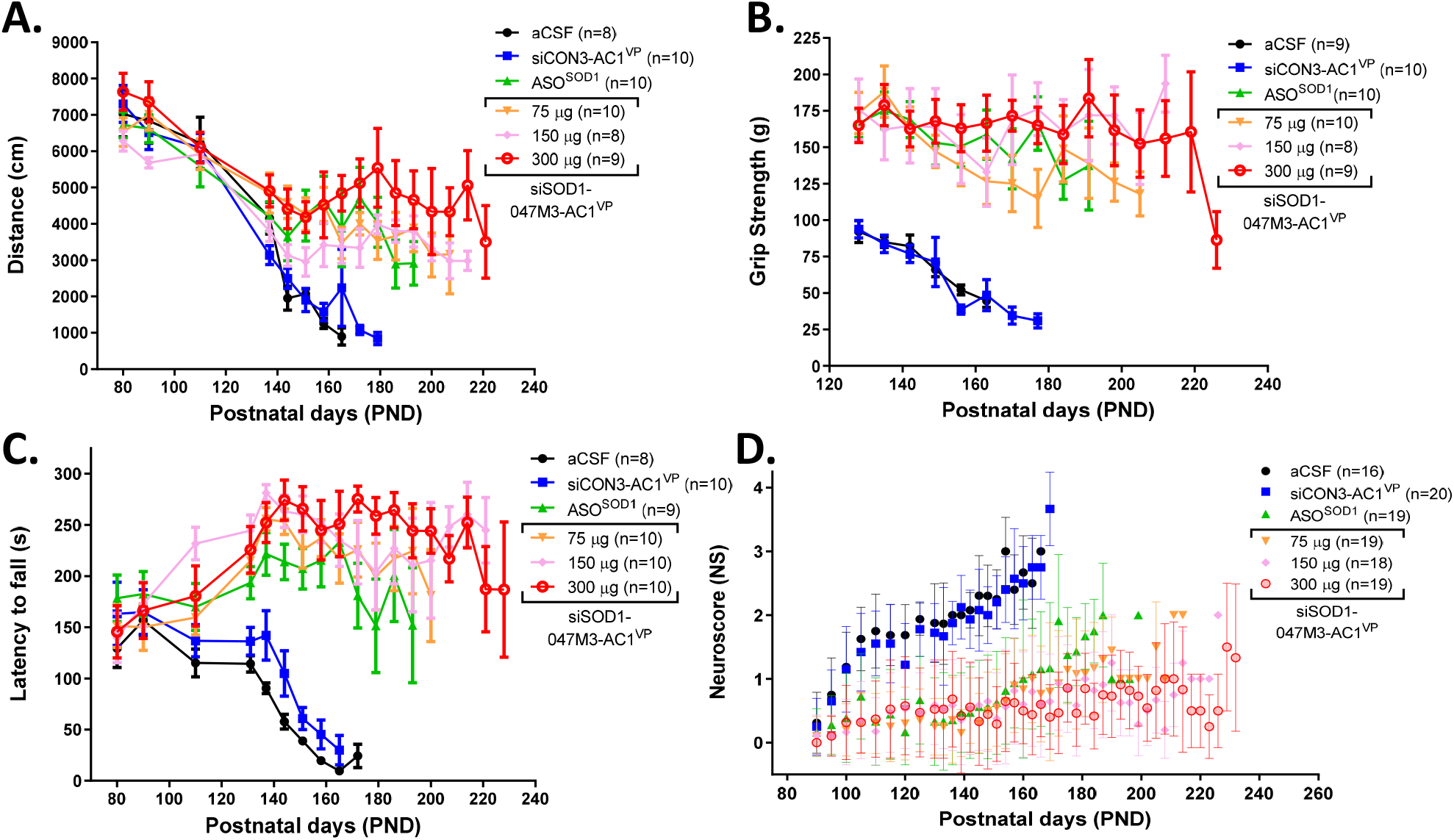
siRNA-ACO treatment improves motor function in SOD1^G93A^ mice. **A**. Male and female hSOD1^G93A^ mice treated twice with the indicated doses were subject to open field tests during the daytime. Total distance each animal moved was autonomously recorded in centimeters (cm) over the course of 15 minutes. Data is plotted as the mean ± SD distanced traveled for each treatment group. **B**. Grip strength was assessed in male and female animals organized by treatment group. Grip strength tests were performed in triplicate and the average value was recorded for each animal. Data is plotted as mean ± SD for grip strength for each treatment group in grams (g). **C**. Animal fatigue and coordination was assessed by rotarod test for 5 minutes. Experiments were performed in triplicate in which the longest latency time to fall was recorded in seconds (s) for each animal. **D**. Motor function was scored using the ALS TDI neuroscore (NS) scale for all animals prior to open field, rotarod, and/or grip strength tests. Mean NS ± SD is shown at each treatment group at the indicated time points.

Motor function was also scored for all animals prior to open field, rotarod, and/or grip strength tests using the ALS Therapy Development Institute (ALS TDI) neuroscore (NS) system. As shown in **Figure 7D**, all mice developed abnormal splay (*i.e*., NS2) by approximately PND130 in both aCSF and siCON3-AC1^VP^ control groups that continued to increase in severity over time (i.e., ≥NS3). Conversely, mean scores for all siSOD1-047M3-AC1^VP^ doses never surpassed NS2 at any time point during the course of the study particularly for the highest dose group (*i.e*., 300 μg) in which mean NS remained predominantly flat at around NS1. ASO^SOD1^ treatment also demonstrated an improved NS comparative to vehicle and siCON3-AC1^VP^ controls. However, by PND150, NS began to increase at a slope similar to controls despite ASO^SOD1^ having ~3X molecular excess than that of siSOD1-047M3-AC1^VP^ at 150 μg dose.

## DISCUSSION

The first clinical study targeting SOD1 for the treatment of ALS was a first-generation ASO called ISIS 333611 (19). While results were presented as proof of concept, first-generation ASOs have had limited clinical success due to a narrow therapeutic index. It was not until the advent of second-generation ASOs that Tofersen (BIIB067) was developed demonstrating improved safety and knockdown activity of SOD1 (6). However, RNAi is still the preferred tool for gene silencing, in part, by being a more potent modality (8, 20). Yet development for indications like ALS have been historically hampered by poor delivery and limited biodistribution of siRNA in target tissues (*i.e*., CNS).

Only recently have new platforms emerged enabling delivery to the CNS including the siRNA-ACO technology disclosed within this study. By screening siRNAs and implementing chemical modification, we identified siRNA drug candidates with low picomolar potency for knockdown of SOD1 in model cell lines. Conjugation to AC1 enabled broad knockdown activity in animal tissues throughout the brain and spinal cord, while maintaining potency comparatively better than a second-generation ASO identical to Tofersen in sequence and chemistry (*i.e*., ASO^SOD1^). Further modification with 5’VP provided enhanced knockdown durability and demonstrated therapeutic efficacy (*i.e*., delayed disease progression, prolonged survival, and improved motor function) in SOD1^G98A^ mice at molar equivalent doses ≥3X lower than ASO^SOD1^.

An added benefit of siRNA-ACO is the simplicity of its manufacturing. Synthesis including conjugation is compatible with classic oligonucleotide solid-phase chemistry in which the entire process is performed on-support. Other than 5’VP, the disclosed siRNA-ACO sequences were comprised entirely of standard amidites (*i.e*., PS, 2’Ome, 2’F, and 2’MOE) with known pre-clinical and clinical safety data. In addition, GMP manufacturing of siRNA-ACO to scale has already been validated via contract manufacturing.

Although ACO conjugation was developed to impart self-delivery properties similar to ASOs, the precise mechanism by which it facilitates siRNA delivery has not been fully elucidated. *In vitro* assays with mouse serum revealed a supershift in gel migration only for the siRNA-ACOs indicative of protein binding. This is reminiscent of the protein interactions that have been reported to influence aspects of ASO performance *in vivo* including biodistribution and cellular uptake (21). While grafting ASO chemistry onto siRNAs has been explored before (*i.e*., asymmetric sd-rxRNA), this is to the best of our knowledge the first example in which conjugation of a short non-targeting oligonucleotide enables robust activity in the CNS (22). In this context, ACO composition is readily amendable to changes in sequence, length, and/or chemistries for further optimization as canonical ASO function via complementarity to a cognate sequence is irrelevant for siRNA-ACO knockdown activity.

Our current lead candidate (*i.e*., siSOD1-047M3-AC1^VP^) represents our first iteration of the siRNA-ACO technology for RNAi delivery to the CNS. With future studies focused on safety pharmacology and toxicology, we aim to clinically validate this delivery platform using ALS as an exemplary indication. However, prevalence of pathogenic mutations located within the siRNA target site of the SOD1 transcript must be considered for estimating actual patient population size. For instance, out of the two pathogenic SNPs located in the target sequence of siSOD1-047M3-AC1^VP^, only P.F21C was partially resilient to knockdown. P.F21C has been linked to patients of Chinese and Korean descent who present with familial ALS and comprise approximately 2.7% of the total ALS^SOD1^ population (18, 23–25). As such, P.F21C may serve as genetic marker for drug efficacy and/or patient exclusion in clinical study design.

## MATERIAL AND METHODS

### High-throughput screening of siRNAs targeting human SOD1

Open reading frame (ORF) from human SOD1 (hSOD1) cDNA sequence (NM_000454.5) served as the template for siRNA design via in-house algorithm. A total of 268 duplexes were synthesized at 19-nucleotides in length without medicinal chemistry (**Supplementary Table 3)**. Plating and transfection of 293A cells was performed in 96-well plates each containing 32 siRNAs and 8 quality control treatments at 2 concentrations (*i.e*., 0.1 and 10 nM) in duplicate. Cells were cultured for 24 hours and lysis was automated via the Fluent System 780 liquid handling system (Tecan, Hombrechtikon, The Switzerland) using an optimized formula containing propidium iodide (PI) based on the Cell-Lysis (CL) buffer for one-step RT-qPCR as previous described (26). Integration of PI in sample preparation served to monitor variation in cell number (*e.g*., untoward cytotoxicity) by staining total nucleic acid content in crude lysates. Staining was quantified via optical density (OD) at 535 nm excitation and 615 nm emission wavelengths on the Infinite M200 Pro microplate reader (Tecan). Samples were subsequently transferred to 384-well plates for analysis by RT-qPCR on the 480 Real-Time PCR system (Roche, Basel, Switzerland) using the One-Step TB Green PrimeScript RT-PCR Kit II (Takara, Kyoto, Japan). Preparation of PCR reactions was automated by the Echo 525 Acoustic Liquid Handler (Beckman Coulter, Brea, CA, USA). A secondary screen was subsequently performed only on the top 30 performing siRNAs at 6 concentrations (*i.e*., 0.0064 - 20 nM). All samples were amplified in triplicate.

### siRNA synthesis

Oligonucleotide sequences were synthesized in-house at Ractigen Therapeutics (Rudong, China) on solid-phase support using a HJ-12 synthesizer (Highgene-Tech Automation, Beijing, China) and subsequently purified via RP-HPLC using an acetonitrile gradient over a UniPS column (NanoMicro Technology, Suzhou, China). Each sequence was reconstituted in sterile water via buffer exchange. Equal molar quantities of each strand were annealed into their corresponding duplexes by briefly heating strand mixtures and cooling to room temperature. Resolution of a single band via gel electrophoresis at predicted molecular weights was used to qualify duplex formation. ESI-MS was used to confirm duplex identity, while overall purity was analyzed via SEC-HPLC using a XBridge Protein BEH SEC 125 A column (Waters Corporation, Milford, MA, USA). Endotoxin levels in each batch were quantified using the end-point Chromogenic Endotoxin Quant Kit (Bioendo, Xiamen, China) via proenzyme Factor C. All control duplexes and chemically modified sequences are listed in **Supplementary Table 4**.

### Cell culture and transfection

293A cells (Cobioer, Nanjing, China, Cat# CBP60436) and SK-N-AS (Procell, Wuhan, China, Cat# CL-0621) cells were maintained in DMEM medium supplemented with 10% FBS, penicillin (100 U/ml) and streptomycin (100 mg/ml). T98G cells (Cobioer, Cat# CBP60301) were maintained in MEM medium supplemented with 10% FBS, 1% NEAA, sodium pyruvate (1 mM), penicillin (100 U/ml) and streptomycin (100 ug/ml). All cell lines were cultured in a humidified atmosphere of 5% CO_2_ at 37°C. Transfections were carried out using Lipofectamine RNAiMax (ThermoFisher, Waltham, MA, USA) in growth media without antibiotics according to the manufacture’s protocol.

### Gene expression analysis via RT-qPCR

Animal tissue frozen in RNALater (Sigma-Aldrich, St. Louis, MO, USA) was homogenized in Total RNA Isolation Reagent (Biosharp, Hefei, China) using a Bioprep-24 Homogenizer (Allsheng, Hangzhou, China). Chloroform was added to the homogenate in which the aqueous phase was removed and mixed with isopropanol. Total RNA was extracted from the tissue prep using the RNeasy RNA kit (Qiagen) according to the manufacture’s protocol. RNA from cell culture was extracted using the Auto-Pure 96A (Allsheng) nucleic acid extraction system. Reverse transcription (RT) reactions were performed with 1 μg total RNA using the PrimeScript RT kit with gDNA Eraser (Takara). The resulting cDNA was amplified in triplicate on the 480 Real-Time PCR system (Roche) using SYBR Premix Ex Taq II (Takara) in conjunction with primer sets specific to human SOD1 (hSOD1) and an internal control for either human (*i.e*., TBP) or mouse (*i.e*., mTbp) samples. Melting curves were performed after amplification to confirm primer specificity. Averaged Ct values for each sample were used to calculate relative gene expression via ΔΔCt method. Primer sequences are available in **Supplementary Table 4**.

### Caspase 3/7 activity assay

Caspase 3/7 activity was quantified in cell culture by using the Caspase-Glo 3/7 assay system (Promega, Madison, WI, USA). Briefly, a luminogenic substrate was added directly to culture media and incubated for 20 mins at 37°C. Luminescence was subsequently measured on an Infinite M200 Pro microplate reader (Tecan). Relative Caspase 3/7 activity was calculated by subtracting background signal of blank from the luminescence values in each well and normalizing data to non-treated (Mock) controls.

### Cell viability assay

*In vitro* cell viability was measured using the CCK-8 assay (Dojindo, Mashiki-machi, Japan) according to the manufacture’s protocol. Briefly, fresh media containing WST-8 substrate was added to each well of the tissue culture plate and incubated for at least 1 hour at 37°C. Absorbance was measured at 450 nm on an Infinite M200 Pro microplate reader (Tecan). Relative viability is calculated by subtracting background absorbance of the blank control from the OD values in each well and normalizing data to non-treated (Mock) controls.

### Gel shift assay

siRNA or siRNA-ACO duplexes were incubated at 37°C in 50% human serum at a final concentration of 3 μM for ≤1hr. Samples were subsequently mixed with 10X Loading Buffer (Takara) and resolved on agarose gels using GelRed Nucleic Acid Gel Stain (Biotium, Fremont, CA, USA) to visualize shifts in gel migration. Gel images were captured on the ChemiDoc XRS+ Imager System (Biorad, Hercules, CA, USA).

### Luciferase reporter constructs and knockdown assessment

Target sequence containing either P.E22G or P.F21C mutant SNPs were cloned into the multiple cloning site (MCS) of luciferase reporter vector pmirGLO (Promega) between the NheI and SalI restriction enzyme (RE) sites downstream of the firefly luciferase gene (luc2) to generate constructs pLuc^P.E22G^ and pLuc^P.F21C^, respectively. A control reporter construct (*i.e*., pLuc^SOD1^) was also created containing consensus hSOD1 sequence perfectly complementary to siSOD1-047M3-AC1^VP^ guide strand. All constructs were subcloned in DH5a bacteria (Tolobio, Shanghai, China) and colonies were selected for DNA sequencing to confirm insertion of target sequence. Exemplary colonies were scaled up for plasmid isolation via midiPrep (Qiagen, Hilden, Germany). 293A cells were plated in 96-well cell culture plates at 30,000 cells/well in absence of antibiotics. Cells were co-transfected with one of reporter plasmids (*i.e*., pLuc^P.E22G^, pLuc^P.F21C^, or pLuc^SOD1^) at 100 ng/well in combination with siSOD1-047M3-AC1^VP^ or scramble control at the indicated concentrations using 0.3 μL of Lipofectamine 2000 (ThermoFisher). Wells treated in absence of test article (0 nM) served as non-treated controls. Cells were cultured for 24 hours and luciferase activity was quantified using the Dual-Glo Luciferase Assay System (Promega) according to the manufacture’s protocol. Briefly, cells were lysed in 50 μL Passive lysis buffer (Promega) in which 20 μL of lysate was mixed with 20 μL of Dual-Glo Luciferase Reagent and incubated for 10 mins at room temperature. Luminescence was subsequently measured on an Infinite 200 Pro microplate reader (Tecan) to quantify luciferase activity. Following measurements, 20 μL of Dual-Glo Stop & Glo Reagent (Promega) was added to each well and incubated for an additional 10 mins at room temperature. Luminescence was again measured to quantify *Renilla* activity, which served to normalize luciferase reporter results. Data was calculated as the ratio of reporter luminescence to *Renilla* luminescence relative to the ratio of non-treated controls. Percent knockdown (% KD) was calculated as 1-(Ratio_siRNA_/Ratio_non-treated_)*100.

### Animal handling and grouping

Parental transgenic hSOD1^G93A^ mice (Strain ID #004435) were purchased from The Jackson Laboratory (Bar Harbor, ME, USA) and imported into China via Nantong University (Nantong City, Jiangsu Province, China). Mice were delivered to the animal facility at 6 weeks of age and subsequently bred domestically at Nantong University who supplied the animals for this study. All animal procedures were approved by the IACUC at Nantong University. Formulations for animal treatments were prepared fresh prior to use by dissolving allotments of lyophilized oligonucleotide into aCSF to create stock solutions for dilution to the intended treatment concentrations. Animals were randomly allocated into study groups based on body weight and sex. Any animals in poor health or with obvious abnormalities were omitted from the experiments. Randomization was analyzed using Ordinary one-way ANOVA via GraphPad Prism version 8.3.0 Windows (GraphPad Software, San Diego, CA, USA). Female hSOD1^G93A^ mice typical weighed approximately 20-25% less than their male liter mates.

### Intracerebroventricular (ICV) injection

Avertin (1.2%) was prepared fresh and sterilizes via 0.2-micron filter. Mice were dosed at 0.30-0.35 ml per 10 g body weight via intraperitoneal (IP) injection in a stereotaxic apparatus to rapidly induce anesthesia for up to 30 minutes. An approximate 11.5 mm incision was made in the animal’s scalp and a 25-gauge needle attached to a Hamilton syringe containing the appropriate siRNA formulation was placed at bregma level. The needle was moved to the appropriate anterior/posterior and medial/lateral coordinates (0.2 mm anterior/posterior and 1 mm to the right medial/lateral). A total of 10 μL was injected into the lateral ventricle at an approximate rate of 1 μl/s. Following treatment, the needle was slowly withdrawn and the wound sutured close.

### Intrathecal (IT) injection

Anesthesia was administered via 3.0% isoflurane in an induction chamber for continuous 10 mins. Hair was shaved around the injection site at the base of the tail and cleaned with 75% ethanol. The space between the L5-L6 spinous processes was identified and a 30-gauge needle attached to a microliter syringe containing the appropriate drug formulations was slowly inserted into the intradural space until a tail flick was observed. The needle position was subsequently secured in which 10 μL total volume of solution was injected over the course of 1 min.

### Quantification of siRNA-ACO in animal tissues

Tissue lysate was prepared in lysis buffer (0.5% CA-630, 1mM EDTA, 150 mM NaCl) using a Bioprep-24 Homogenizer (Allsheng). Samples were subsequently heated to 95°C to inactivate sample proteins. Serial dilution of non-treated lysate spiked with siRNA-ACO was used to generate 8-point standard curves. Reverse transcription (RT) reactions were performed using the PrimeScript RT reagent kit (Takara) in conjugation with custom stem-loop primers specific to siRNA guide strands. Each sample was amplified in triplicate on the 480 Real-Time PCR system (Roche) using SYBR Premix Ex Taq II (Takara) reaction mix with primer sets specific to guide strand cDNA. Melting curves were performed after amplification to confirm primer specificity. Absolute quantities of siRNA were extrapolated by linear regression using the appropriate standard curves. Tissue concentrations were calculated as the ratio of absolute siRNA mass (ng) relative to the total weight (g) of tissue sample prepped for lysis.

### Clinical observation and endpoint criteria

Animals were observed after injection for up to 4 hours and daily thereafter until endpoint. Body weight was determined before test substance administration and at recorded intervals thereafter. Animals with weight loss >20% relative to their initial mass at day of treatment or a neuroscore of NS4 met endpoint criteria.

### Neurological scoring

For animals treated via IT injection, mice were evaluated for signs of motor deficit using the ALS Therapy Development Institute (ALS TDI) neuroscore (NS) system, which was developed to provide unbiased assessment of disease progression based on hindlimb dysfunction common to SOD1^G93A^ mice (27). NS was assigned based on the following 4-point scale: 0 if no signs of motor dysfunction (*i.e*., pre-symptomatic), 1 if hindlimb tremors are evident when suspended by tail (*i.e*., first symptoms), 2 if gait abnormalities are present (*i.e*., onset of paresis), 3 if dragging at least 1 hindlimb (*i.e*., partial paralysis), and 4 if inability to right itself within 10 seconds (*i.e*., endpoint paralysis).

### Open field test

Each mouse was placed in the corner of an open field apparatus (50 cm length × 50 cm width × 50 cm height) during daylight hours and allowed to freely roam for 15 minutes. An overhead camera recorded travel path of each animal. Video footage was analyzed by automated tracking software Samart 3.0 (Bioseb, Vitrolles, France) to calculate total distance traveled.

### Rotarod analysis

Animals were trained for 3 days prior to data acquisition. Mice were placed on a motionless rotarod apparatus (XinRuan Information Technology, Shanghai, China) with a swivel bar 60 mm in diameter. Rotational speed was accelerated from 0-30 rpm over the course of 300 seconds. Latency time was recorded as the amount of time it took for each animal to fall off the swivel bar. Each animal was tested in triplicate in which the longest value represents latency time.

### Grip strength test

Mice were lowered onto a grid plate in which its forepaws and hind paws were allowed to grasp the grid. The tail was gently pulled and maximal muscle strength was measured on the XR501 grip strength meter (XinRuan Information Technology) in units of mass until the animal relinquished its grasp. Each animal was tested in triplicate in which the mean value represents grip strength.

### Statistical analysis

Data analytics were performed using GraphPad Prism version 8.3.0 Windows. Dose response curves and IC_50_ values were extrapolated using non-linear regression via 4-parameter concentration-inhibition model. Where specified, mean values were compared using Tukey’s multiple comparison test to determine statistical differences between different dose response curves. Drug quantities in tissue in relationship to knockdown activity including extrapolation of ED_50_ values were performed using non-linear regression via 3-parameter concentration-response model. Time-stratified data (*i.e*., peak weight analysis and animal survival) was plotted via Kaplan-Meier graphs in which statistical significance was verified using the Mantel-Cox test.

