## Supplementary Figures 1-10 for "Local administration of a novel siRNA modality into the CNS extends survival and improves motor function in the SOD1^G93A^ mouse model for ALS"

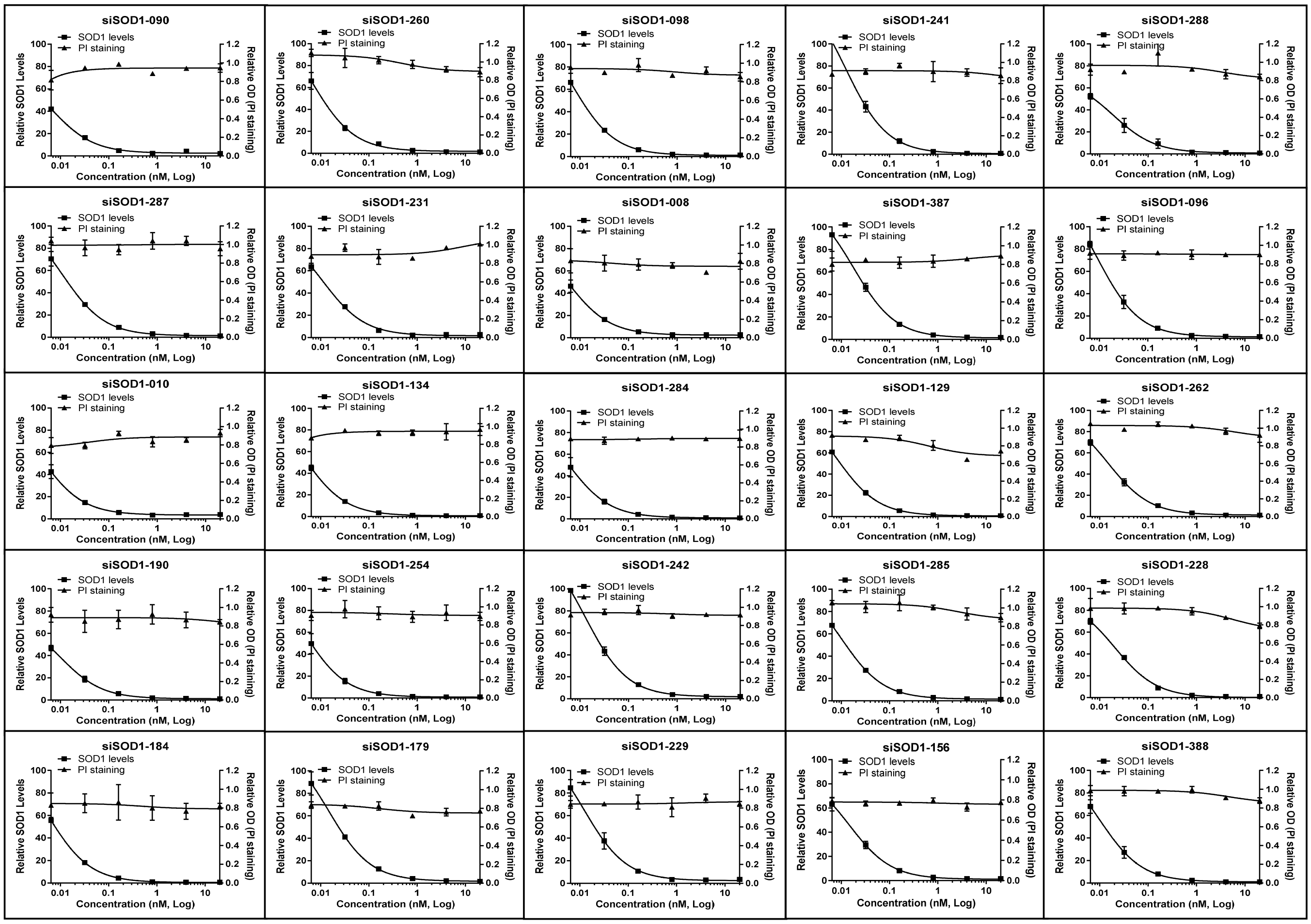


**Supplementary Figure 1. Secondary screening of siRNA potency and untoward cytotoxicity *in vitro*.** Knockdown activity and cell viability in 293A cells of the remaining top 25 performing siRNAs identified in the initial screen is shown following transfection at 6 escalating concentrations (*i.e.,* 0.0064, 0.032, 0.16, 0.8, 4, and 20 nM). SOD1 knockdown and untoward cytotoxicity were quantified by RT-qPCR and PI staining relative to Mock treatments, respectively.

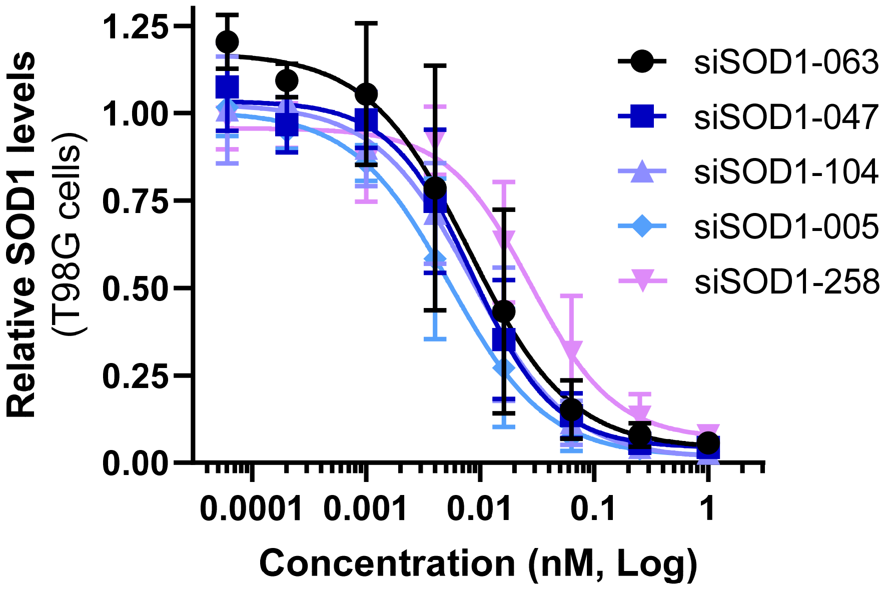


**Supplementary Figure 2. Dose-dependent knockdown of the top 5 siRNA candidates in T98G cells.** Dose response curves were generated in T98G cells following transfection for 24 hours with each of the top 5 siRNAs at 8 treatment concentrations (*i.e.*, 0.00006, 0.0002, 0.001, 0.004, 0.016, 0.063, 0.25, and 1 nM). SOD1 expression levels were quantified via RT-qPCR using gene specific primer sets. TBP was amplified as an internal reference used to normalize expression data. Shown are mean values ± SD from 2 experimental replicates relative to controls treated in absence of oligonucleotide.

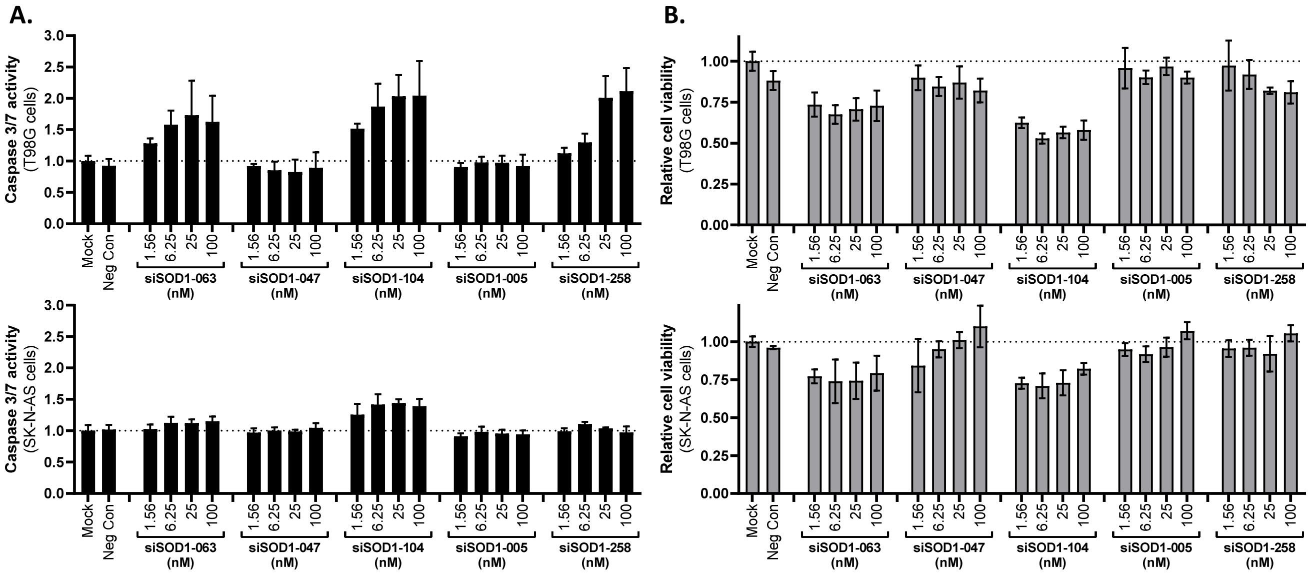


**Supplementary Figure 3. Untoward cytotoxicity of the top 5 siRNA candidates *in vitro*. A-B.** SK-N-AS and T98G cells were transfected with each of the top 5 siRNAs at 4 treatment concentrations in noted excess of their $IC_{50}$ values for SOD1 knockdown (*i.e.*, 1.56, 6.25, 25, and 100 nM). Treatment with siCon at 100 nM served as a negative control (Neg Con) for untoward cytotoxicity. Mock samples were transfected in absence oligonucleotide. Untoward cytotoxicity was evaluated 72 hours after treatment by quantifying both caspase 3/7 activity (**A**) and metabolism of WST-8 reagent as a marker for cell viability (**B**). Data is shown as mean values ± SD from 2 experimental replicates relative to Mock treatments (dotted lines).

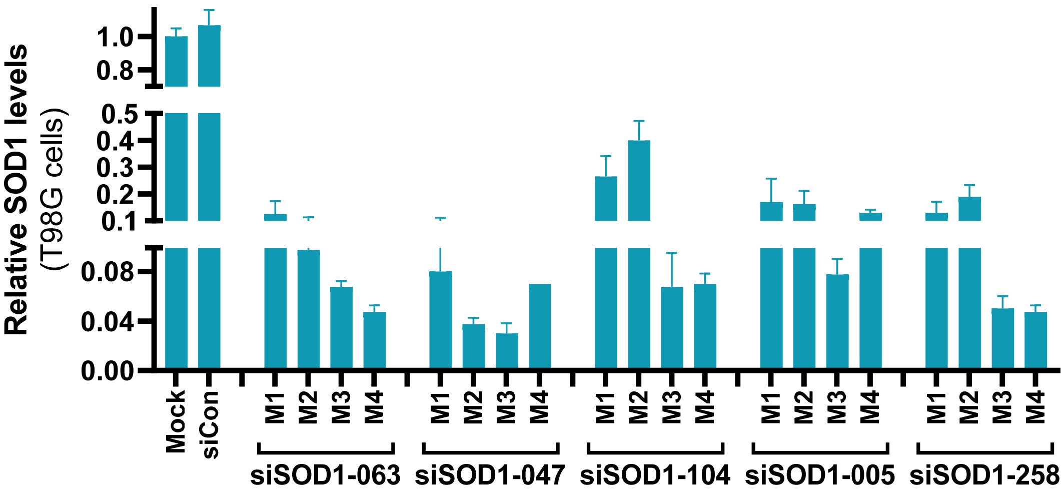


**Supplementary Figure 4. Impact of different chemical modifications patterns on siRNA knockdown activity in T98G cells.** T98G cells were transfected at 0.1 nM with 4 different chemically modified variants of each siRNA candidate (*i.e.*, M1, M2, M3, or M4) or a non-specific siRNA control (siCon) for 24 hours. Knockdown activity was assessed via RT-qPCR relative to Mock treatment. Data represents mean ± SD from 2 experimental replicates.

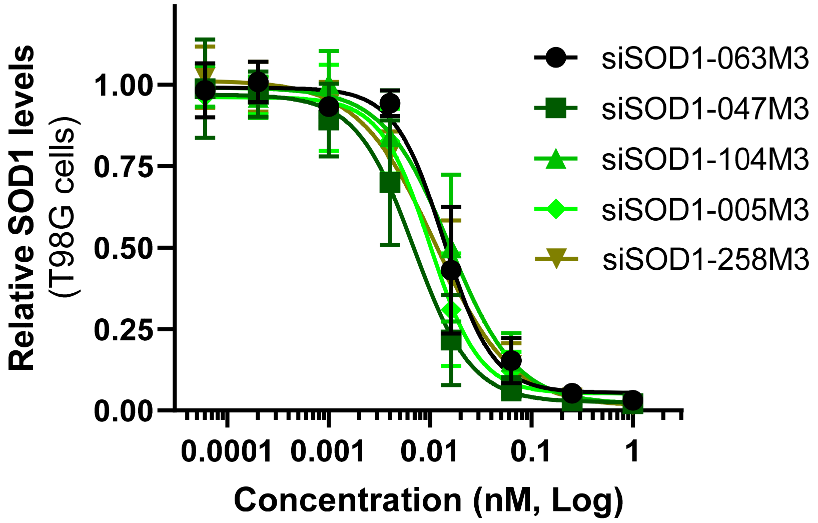


**Supplementary Figure 5. Dose-dependent knockdown of M3-modified siRNAs in T98G cells.** Dose response curves were generated in T98G cells following transfection for 24 hours with each M3-modified siRNA candidate (*i.e.*, siSOD1-063M3, 047M3, 104M3, 005M3, and 258M3) at 8 treatment concentrations (*i.e.*, 0.00006, 0.0002, 0.001, 0.004, 0.016, 0.063, 0.25, and 1 nM). SOD1 expression levels were quantified via RT-qPCR using gene specific primer sets. TBP was amplified as an internal reference used to normalize expression data. Shown are mean values ± SD from 3 experimental replicates relative to samples treated in absence of oligonucleotide.

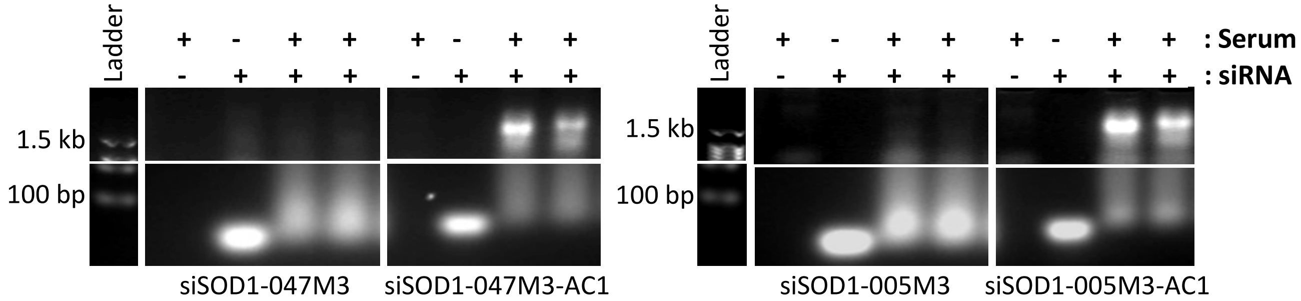


**Supplementary Figure 6. ACO conjugation to chemically modified siRNA promotes novel interactions in human serum.** Chemically modified siRNAs (*i.e.*, siSOD1-047M3 and siSOD1-005M3) or ACO conjugates (*i.e.*, siSOD1-047M3-AC1 and siSOD1-005M3-AC1) were incubated with human serum for ≤1 hour. Samples were resolved on agarose gels to visualize shifts in siRNA migration. Shown are gel images stained for nucleic acid content including DNA ladder (Ladder) for reference. Binding reactions are shown in duplicate. Serum alone served as a control for background signal. Duplexes incubated in absence serum define migration of unbound siRNA.

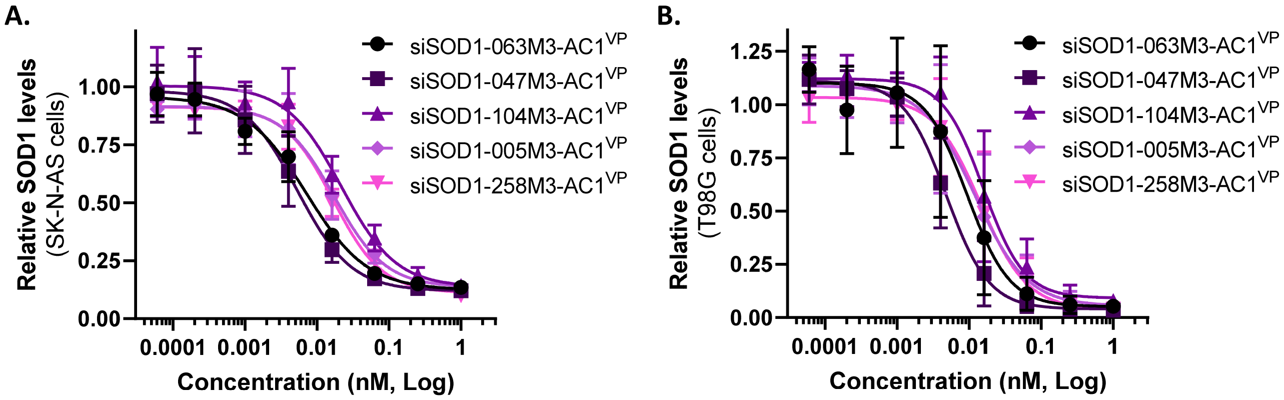


**Supplementary Figure 7. Knockdown activity of siRNA-ACO drug candidates *in vitro*. A-B.** Dose response curves were generated in SK-N-AS (**A**) and T98G (**B**) cells for each 5'VP-modified siRNA-ACO (*i.e.*, siSOD1-063M3-AC1$^{VP}$, 047M3-AC1$^{VP}$, 104M3-AC1$^{VP}$, 005M3-AC1$^{VP}$, and 258M3-AC1$^{VP}$) at 8 treatment concentrations (*i.e.*, 0.00006, 0.0002, 0.001, 0.004, 0.016, 0.063, 0.25, and 1 nM). SOD1 expression levels were quantified via RT-qPCR using gene specific primer sets. TBP was amplified as an internal reference to normalize data. Shown are the expression values of SOD1 mRNA relative to mock transfections. Data represents mean ± SD from 2 experimental replicates.

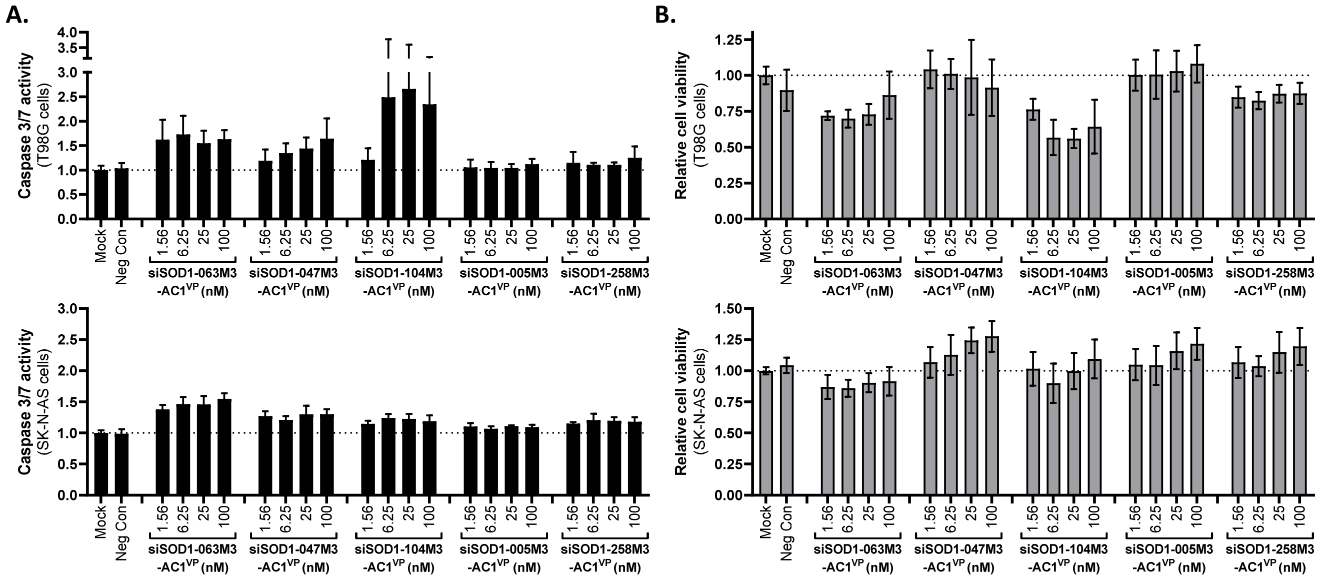


**Supplementary Figure 8. Untoward cytotoxicity of siRNA-ACO drug candidates *in vitro*. A-B.** SK-N-AS and T98G cells were transfected with the lead siRNA-ACO conjugates (*i.e.*, siSOD1-063M3-AC1$^{VP}$, 047M3-AC1$^{VP}$, 104M3-AC1$^{VP}$, 005M3-AC1$^{VP}$, and 258M3-AC1$^{VP}$) at 4 treatment concentrations in excess of their $IC_{50}$ values (*i.e.*, 1.56, 6.25, 25, and 100 nM). Treatment with siCon at 100 nM served as a negative control (Neg Con) for untoward cytotoxicity. Mock samples were transfected in absence oligonucleotide. Untoward cytotoxicity was evaluated 72 hours after treatment by quantifying both caspase 3/7 activity (**A**) and metabolism of WST-8 reagent as a marker for cell viability (**B**). Data is shown as mean values ± SD from 2 experimental replicates relative to Mock treatments (dotted lines).

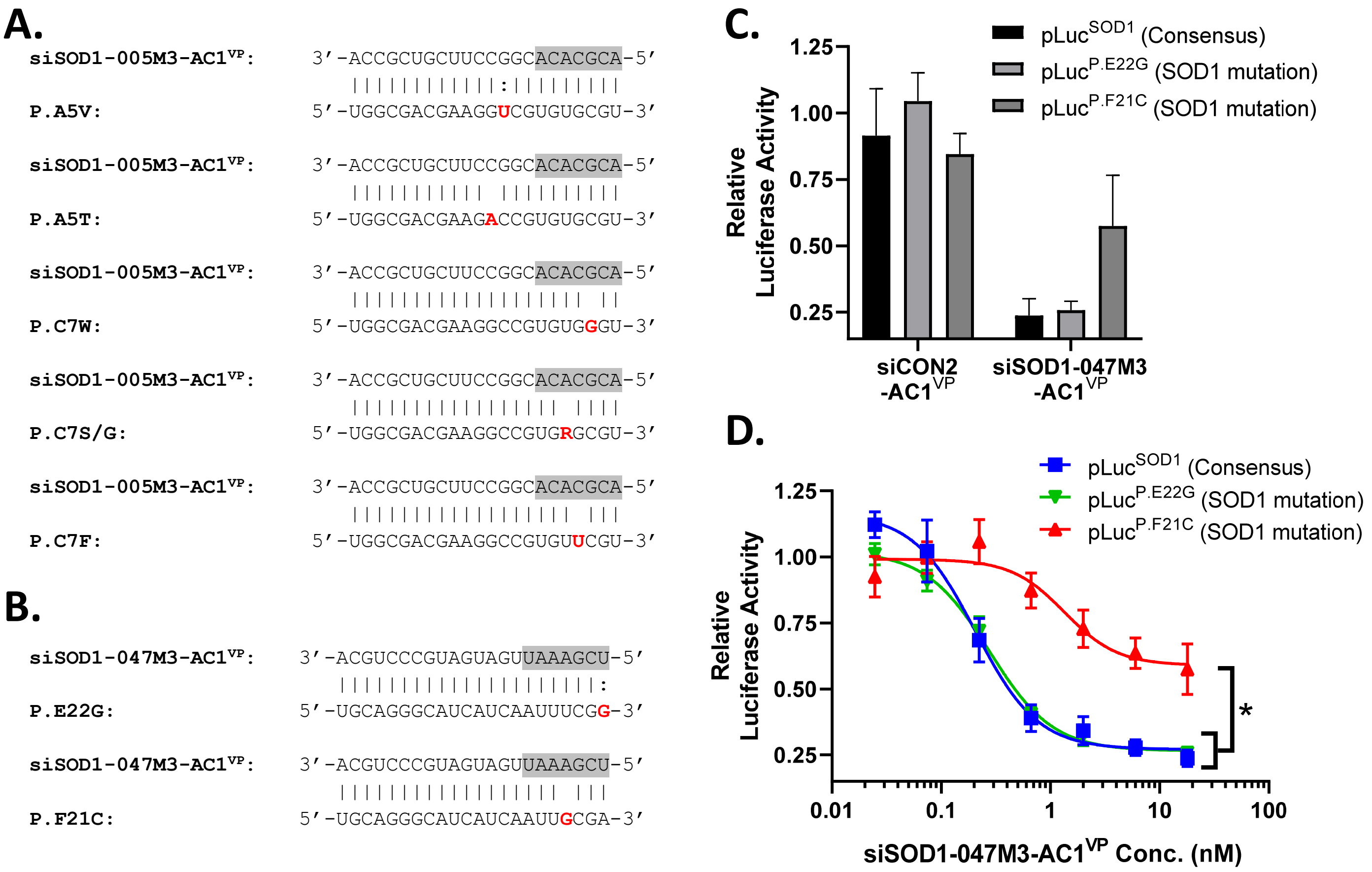


**Supplementary Figure 9. Pathogenic SNPs in target sites of lead siRNA-ACO candidates. A-B.** Location of pathogenic SNPs within the target sites of siSOD1-005M3-AC1$^{VP}$ (**A**) and siSOD1-047M3-AC1$^{VP}$ (**B**). Shown is guide strand sequence including "seed" region (highlighted in grey) complementary to target sites in hSOD1 transcript containing the indicated pathogenic SNPs. Nucleotide mutation is shown in red in which 'R' signifies a purine substitution. **C.** Luciferase reporter constructs (*i.e.*, pLuc$^{SOD1}$, pLuc$^{P.E22G}$, pLuc$^{P.F21C}$) containing either consensus sequence or one of the pathogenic mutations (*i.e.*, P.E22G and P.F21C) were co-transfected with siSOD1-047M3-AC1$^{VP}$ or a non-specific scramble control (siCON2-AC1$^{VP}$) at 18 nM concentrations in 293A cells. **D.** Dose response curves for luciferase activity were generated following co-transfection with siSOD1-047M3-AC1$^{VP}$ at escalating concentrations (*i.e.*, 0.03, 0.07, 0.22, 0.67, 2.0, 6.0, and 18 nM). Data represents mean ± SEM from 2 experimental replicates relative to samples treated in absence of siRNA. Statistical significance (*) was determined using Tukey's multiple comparison test to compare the mean values at each data point within the three dose response curves.

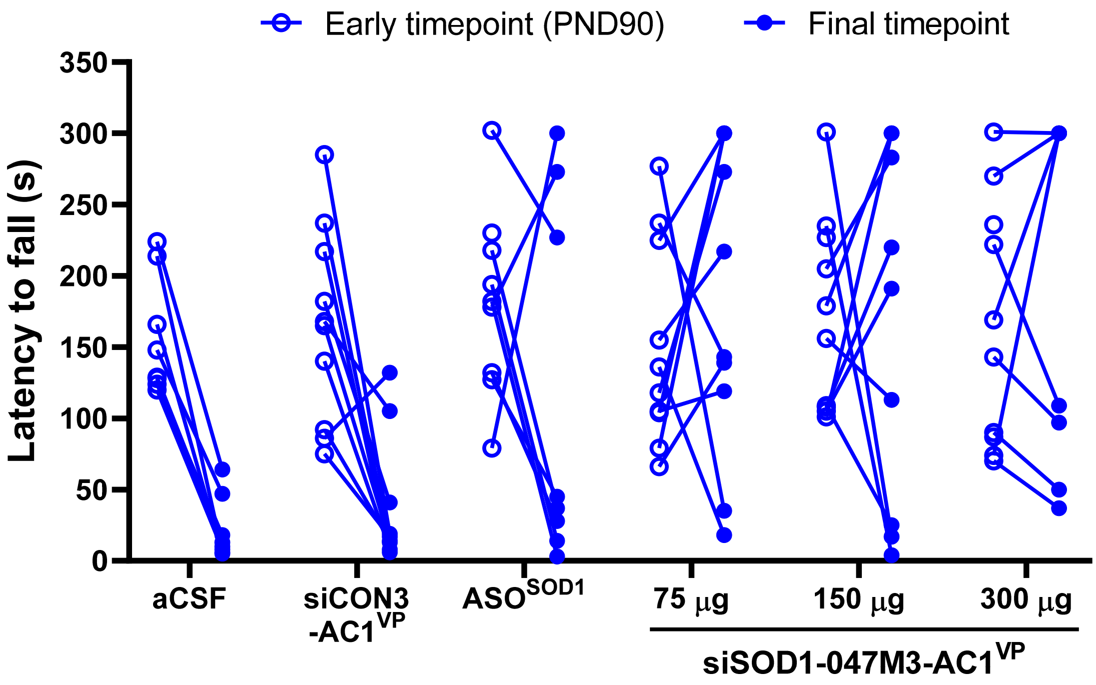


**Supplementary Figure 10. Temporal comparison of rotarod performance for each individual animal following siRNA-ACO treatment.** Latency time to fall in seconds (s) as measured by the rotarod test for each individual animal in their corresponding treatment groups at an early timepoint (*i.e.*, PND90) in comparison to performance at end-stage (*i.e.*, final timepoint). All tests were performed in triplicate in which the longest latency time was recorded for each animal.
