## Supplementary Tables 1-4 for "Local administration of a novel siRNA modality into the CNS extends survival and improves motor function in the SOD1^G93A^ mouse model for ALS"

**Supplementary Table 1.** *In vitro* potency of drug candidates**.**

| **siRNA** | **$IC_{50}$ (pM)*** | |
|---|---|---|
| | **SK-N-AS** | **T98G** |
| siSOD1-063 | **4.7 ± 1.4** | **7.9 ± 6.1** |
| siSOD1-047 | **5.1 ± 1.3** | **8.5 ± 3.6** |
| siSOD1-104 | **5.3 ± 1.7** | **8.5 ± 3.9** |
| siSOD1-005 | **3.4 ± 1.1** | **5.4 ± 2.5** |
| siSOD1-258 | **19.7 ± 10.0** | **27.3 ± 15.7** |
| siSOD1-063M3 | **9.3 ± 3.4** | **13.6 ± 2.9** |
| siSOD1-047M3 | **4.9 ± 1.5** | **6.9 ± 1.9** |
| siSOD1-104M3 | **18.8 ± 7.6** | **15.5 ± 4.8** |
| siSOD1-005M3 | **10.4 ± 7.8** | **9.9 ± 2.2** |
| siSOD1-258M3 | **17.8 ± 7.7** | **11.2 ± 3.1** |
| siSOD1-063M3-AC1$^{VP}$ | **7.1 ± 2.3** | **9.4 ± 6.3** |
| siSOD1-047M3-AC1$^{VP}$ | **5.3 ± 2.5** | **4.8 ± 1.5** |
| siSOD1-104M3-AC1$^{VP}$ | **21.6 ± 10.9** | **15.68 ± 7.1** |
| siSOD1-005M3-AC1$^{VP}$ | **17.3 ± 6.6** | **12.1 ± 7.7** |
| siSOD1-258M3-AC1$^{VP}$ | **16.7 ± 3.7** | **14.3 ± 7.1** |

* $IC_{50}$ vales ± SD

**Supplementary Table 2.** $ED_{50}$ and tissue concentrations of siRNA-ACO to trigger median response**.**

| siRNA-ACO | CNS Tissue | $ED_{50}$ (μg) | Conc. (μg/g)* |
|---|---|---|---|
| **siSOD1-047M3-AC1$^{VP}$** | Cerebellum | 99.8 | 1.22 |
| | Cerebrum | 80.9 | 0.174 |
| | Spinal Cord | 151 | 0.082 |
| **siSOD1-005M3-AC1$^{VP}$** | Cerebellum | 38.8 | *ND* |
| | Cerebrum | 26.4 | 0.079 |
| | Spinal Cord | 25 | 0.045 |

* mass (μg) of siRNA-ACO per gram (g) of tissue.

*ND:* could not be determined

**Supplementary Table 3.** siRNA sequences tested in HTS for SOD1 knockdown.

| siRNA | Sense (5'-3') | Antisense (5'-3') |
|---|---|---|
| **siSOD1-005** | CGACGAAGGCCGUGUGCGUTT | ACGCACACGGCCUUCGUCGTT |
| **siSOD1-008** | CGAAGGCCGUGUGCGUGCUTT | AGCACGCACACGGCCUUCGTT |
| **siSOD1-010** | AAGGCCGUGUGCGUGCUGATT | UCAGCACGCACACGGCCUUTT |
| **siSOD1-011** | AGGCCGUGUGCGUGCUGAATT | UUCAGCACGCACACGGCCUTT |
| **siSOD1-017** | UGUGCGUGCUGAAGGGCGATT | UCGCCCUUCAGCACGCACATT |
| **siSOD1-035** | ACGGCCCAGUGCAGGGCAUTT | AUGCCCUGCACUGGGCCGUTT |
| **siSOD1-037** | GGCCCAGUGCAGGGCAUCATT | UGAUGCCCUGCACUGGGCCTT |
| **siSOD1-038** | GCCCAGUGCAGGGCAUCAUTT | AUGAUGCCCUGCACUGGGCTT |
| **siSOD1-040** | CCAGUGCAGGGCAUCAUCATT | UGAUGAUGCCCUGCACUGGTT |
| **siSOD1-041** | CAGUGCAGGGCAUCAUCAATT | UUGAUGAUGCCCUGCACUGTT |
| **siSOD1-042** | AGUGCAGGGCAUCAUCAAUTT | AUUGAUGAUGCCCUGCACUTT |
| **siSOD1-043** | GUGCAGGGCAUCAUCAAUUTT | AAUUGAUGAUGCCCUGCACTT |
| **siSOD1-044** | UGCAGGGCAUCAUCAAUUUTT | AAAUUGAUGAUGCCCUGCATT |
| **siSOD1-045** | GCAGGGCAUCAUCAAUUUCTT | GAAAUUGAUGAUGCCCUGCTT |
| **siSOD1-046** | CAGGGCAUCAUCAAUUUCGTT | CGAAAUUGAUGAUGCCCUGTT |
| **siSOD1-047** | AGGGCAUCAUCAAUUUCGATT | UCGAAAUUGAUGAUGCCCUTT |
| **siSOD1-050** | GCAUCAUCAAUUUCGAGCATT | UGCUCGAAAUUGAUGAUGCTT |
| **siSOD1-051** | CAUCAUCAAUUUCGAGCAGTT | CUGCUCGAAAUUGAUGAUGTT |
| **siSOD1-052** | AUCAUCAAUUUCGAGCAGATT | UCUGCUCGAAAUUGAUGAUTT |
| **siSOD1-053** | UCAUCAAUUUCGAGCAGAATT | UUCUGCUCGAAAUUGAUGATT |
| **siSOD1-056** | UCAAUUUCGAGCAGAAGGATT | UCCUUCUGCUCGAAAUUGATT |
| **siSOD1-057** | CAAUUUCGAGCAGAAGGAATT | UUCCUUCUGCUCGAAAUUGTT |
| **siSOD1-058** | AAUUUCGAGCAGAAGGAAATT | UUUCCUUCUGCUCGAAAUUTT |
| **siSOD1-059** | AUUUCGAGCAGAAGGAAAGTT | CUUUCCUUCUGCUCGAAAUTT |
| **siSOD1-060** | UUUCGAGCAGAAGGAAAGUTT | ACUUUCCUUCUGCUCGAAATT |
| **siSOD1-061** | UUCGAGCAGAAGGAAAGUATT | UACUUUCCUUCUGCUCGAATT |
| **siSOD1-062** | UCGAGCAGAAGGAAAGUAATT | UUACUUUCCUUCUGCUCGATT |
| **siSOD1-063** | CGAGCAGAAGGAAAGUAAUTT | AUUACUUUCCUUCUGCUCGTT |
| **siSOD1-064** | GAGCAGAAGGAAAGUAAUGTT | CAUUACUUUCCUUCUGCUCTT |
| **siSOD1-066** | GCAGAAGGAAAGUAAUGGATT | UCCAUUACUUUCCUUCUGCTT |
| **siSOD1-069** | GAAGGAAAGUAAUGGACCATT | UGGUCCAUUACUUUCCUUCTT |
| **siSOD1-072** | GGAAAGUAAUGGACCAGUGTT | CACUGGUCCAUUACUUUCCTT |
| **siSOD1-073** | GAAAGUAAUGGACCAGUGATT | UCACUGGUCCAUUACUUUCTT |
| **siSOD1-074** | AAAGUAAUGGACCAGUGAATT | UUCACUGGUCCAUUACUUUTT |
| **siSOD1-075** | AAGUAAUGGACCAGUGAAGTT | CUUCACUGGUCCAUUACUUTT |
| **siSOD1-077** | GUAAUGGACCAGUGAAGGUTT | ACCUUCACUGGUCCAUUACTT |
| **siSOD1-079** | AAUGGACCAGUGAAGGUGUTT | ACACCUUCACUGGUCCAUUTT |
| **siSOD1-080** | AUGGACCAGUGAAGGUGUGTT | CACACCUUCACUGGUCCAUTT |
| **siSOD1-085** | CCAGUGAAGGUGUGGGGAATT | UUCCCCACACCUUCACUGGTT |
| **siSOD1-088** | GUGAAGGUGUGGGGAAGCATT | UGCUUCCCCACACCUUCACTT |
| **siSOD1-089** | UGAAGGUGUGGGGAAGCAUTT | AUGCUUCCCCACACCUUCATT |
| **siSOD1-090** | GAAGGUGUGGGGAAGCAUUTT | AAUGCUUCCCCACACCUUCTT |
| **siSOD1-091** | AAGGUGUGGGGAAGCAUUATT | UAAUGCUUCCCCACACCUUTT |
| **siSOD1-092** | AGGUGUGGGGAAGCAUUAATT | UUAAUGCUUCCCCACACCUTT |

| | | |
|---|---|---|
| **siSOD1-093** | GGUGUGGGGAAGCAUUAAATT | UUUAAUGCUUCCCCACACCTT |
| **siSOD1-094** | GUGUGGGGAAGCAUUAAAGTT | CUUUAAUGCUUCCCCACACTT |
| **siSOD1-095** | UGUGGGGAAGCAUUAAAGGTT | CCUUUAAUGCUUCCCCACATT |
| **siSOD1-096** | GUGGGGAAGCAUUAAAGGATT | UCCUUUAAUGCUUCCCCACTT |
| **siSOD1-098** | GGGGAAGCAUUAAAGGACUTT | AGUCCUUUAAUGCUUCCCCTT |
| **siSOD1-099** | GGGAAGCAUUAAAGGACUGTT | CAGUCCUUUAAUGCUUCCCTT |
| **siSOD1-100** | GGAAGCAUUAAAGGACUGATT | UCAGUCCUUUAAUGCUUCCTT |
| **siSOD1-102** | AAGCAUUAAAGGACUGACUTT | AGUCAGUCCUUUAAUGCUUTT |
| **siSOD1-104** | GCAUUAAAGGACUGACUGATT | UCAGUCAGUCCUUUAAUGCTT |
| **siSOD1-105** | CAUUAAAGGACUGACUGAATT | UUCAGUCAGUCCUUUAAUGTT |
| **siSOD1-106** | AUUAAAGGACUGACUGAAGTT | CUUCAGUCAGUCCUUUAAUTT |
| **siSOD1-107** | UUAAAGGACUGACUGAAGGTT | CCUUCAGUCAGUCCUUUAATT |
| **siSOD1-108** | UAAAGGACUGACUGAAGGCTT | GCCUUCAGUCAGUCCUUUATT |
| **siSOD1-114** | ACUGACUGAAGGCCUGCAUTT | AUGCAGGCCUUCAGUCAGUTT |
| **siSOD1-117** | GACUGAAGGCCUGCAUGGATT | UCCAUGCAGGCCUUCAGUCTT |
| **siSOD1-118** | ACUGAAGGCCUGCAUGGAUTT | AUCCAUGCAGGCCUUCAGUTT |
| **siSOD1-119** | CUGAAGGCCUGCAUGGAUUTT | AAUCCAUGCAGGCCUUCAGTT |
| **siSOD1-120** | UGAAGGCCUGCAUGGAUUCTT | GAAUCCAUGCAGGCCUUCATT |
| **siSOD1-122** | AAGGCCUGCAUGGAUUCCATT | UGGAAUCCAUGCAGGCCUUTT |
| **siSOD1-123** | AGGCCUGCAUGGAUUCCAUTT | AUGGAAUCCAUGCAGGCCUTT |
| **siSOD1-125** | GCCUGCAUGGAUUCCAUGUTT | ACAUGGAAUCCAUGCAGGCTT |
| **siSOD1-126** | CCUGCAUGGAUUCCAUGUUTT | AACAUGGAAUCCAUGCAGGTT |
| **siSOD1-127** | CUGCAUGGAUUCCAUGUUCTT | GAACAUGGAAUCCAUGCAGTT |
| **siSOD1-128** | UGCAUGGAUUCCAUGUUCATT | UGAACAUGGAAUCCAUGCATT |
| **siSOD1-129** | GCAUGGAUUCCAUGUUCAUTT | AUGAACAUGGAAUCCAUGCTT |
| **siSOD1-131** | AUGGAUUCCAUGUUCAUGATT | UCAUGAACAUGGAAUCCAUTT |
| **siSOD1-132** | UGGAUUCCAUGUUCAUGAGTT | CUCAUGAACAUGGAAUCCATT |
| **siSOD1-133** | GGAUUCCAUGUUCAUGAGUTT | ACUCAUGAACAUGGAAUCCTT |
| **siSOD1-134** | GAUUCCAUGUUCAUGAGUUTT | AACUCAUGAACAUGGAAUCTT |
| **siSOD1-137** | UCCAUGUUCAUGAGUUUGGTT | CCAAACUCAUGAACAUGGATT |
| **siSOD1-138** | CCAUGUUCAUGAGUUUGGATT | UCCAAACUCAUGAACAUGGTT |
| **siSOD1-140** | AUGUUCAUGAGUUUGGAGATT | UCUCCAAACUCAUGAACAUTT |
| **siSOD1-141** | UGUUCAUGAGUUUGGAGAUTT | AUCUCCAAACUCAUGAACATT |
| **siSOD1-142** | GUUCAUGAGUUUGGAGAUATT | UAUCUCCAAACUCAUGAACTT |
| **siSOD1-148** | GAGUUUGGAGAUAAUACAGTT | CUGUAUUAUCUCCAAACUCTT |
| **siSOD1-150** | GUUUGGAGAUAAUACAGCATT | UGCUGUAUUAUCUCCAAACTT |
| **siSOD1-154** | GGAGAUAAUACAGCAGGCUTT | AGCCUGCUGUAUUAUCUCCTT |
| **siSOD1-156** | AGAUAAUACAGCAGGCUGUTT | ACAGCCUGCUGUAUUAUCUTT |
| **siSOD1-157** | GAUAAUACAGCAGGCUGUATT | UACAGCCUGCUGUAUUAUCTT |
| **siSOD1-158** | AUAAUACAGCAGGCUGUACTT | GUACAGCCUGCUGUAUUAUTT |
| **siSOD1-159** | UAAUACAGCAGGCUGUACCTT | GGUACAGCCUGCUGUAUUATT |
| **siSOD1-160** | AAUACAGCAGGCUGUACCATT | UGGUACAGCCUGCUGUAUUTT |
| **siSOD1-162** | UACAGCAGGCUGUACCAGUTT | ACUGGUACAGCCUGCUGUATT |
| **siSOD1-163** | ACAGCAGGCUGUACCAGUGTT | CACUGGUACAGCCUGCUGUTT |
| **siSOD1-165** | AGCAGGCUGUACCAGUGCATT | UGCACUGGUACAGCCUGCUTT |
| **siSOD1-171** | CUGUACCAGUGCAGGUCCUTT | AGGACCUGCACUGGUACAGTT |
| **siSOD1-173** | GUACCAGUGCAGGUCCUCATT | UGAGGACCUGCACUGGUACTT |

| | | |
|---|---|---|
| **siSOD1-175** | ACCAGUGCAGGUCCUCACUTT | AGUGAGGACCUGCACUGGUTT |
| **siSOD1-176** | CCAGUGCAGGUCCUCACUUTT | AAGUGAGGACCUGCACUGGTT |
| **siSOD1-177** | CAGUGCAGGUCCUCACUUUTT | AAAGUGAGGACCUGCACUGTT |
| **siSOD1-178** | AGUGCAGGUCCUCACUUUATT | UAAAGUGAGGACCUGCACUTT |
| **siSOD1-179** | GUGCAGGUCCUCACUUUAATT | UUAAAGUGAGGACCUGCACTT |
| **siSOD1-180** | UGCAGGUCCUCACUUUAAUTT | AUUAAAGUGAGGACCUGCATT |
| **siSOD1-181** | GCAGGUCCUCACUUUAAUCTT | GAUUAAAGUGAGGACCUGCTT |
| **siSOD1-182** | CAGGUCCUCACUUUAAUCCTT | GGAUUAAAGUGAGGACCUGTT |
| **siSOD1-183** | AGGUCCUCACUUUAAUCCUTT | AGGAUUAAAGUGAGGACCUTT |
| **siSOD1-184** | GGUCCUCACUUUAAUCCUCTT | GAGGAUUAAAGUGAGGACCTT |
| **siSOD1-185** | GUCCUCACUUUAAUCCUCUTT | AGAGGAUUAAAGUGAGGACTT |
| **siSOD1-186** | UCCUCACUUUAAUCCUCUATT | UAGAGGAUUAAAGUGAGGATT |
| **siSOD1-187** | CCUCACUUUAAUCCUCUAUTT | AUAGAGGAUUAAAGUGAGGTT |
| **siSOD1-188** | CUCACUUUAAUCCUCUAUCTT | GAUAGAGGAUUAAAGUGAGTT |
| **siSOD1-189** | UCACUUUAAUCCUCUAUCCTT | GGAUAGAGGAUUAAAGUGATT |
| **siSOD1-190** | CACUUUAAUCCUCUAUCCATT | UGGAUAGAGGAUUAAAGUGTT |
| **siSOD1-192** | CUUUAAUCCUCUAUCCAGATT | UCUGGAUAGAGGAUUAAAGTT |
| **siSOD1-196** | AAUCCUCUAUCCAGAAAACTT | GUUUUCUGGAUAGAGGAUUTT |
| **siSOD1-197** | AUCCUCUAUCCAGAAAACATT | UGUUUUCUGGAUAGAGGAUTT |
| **siSOD1-198** | UCCUCUAUCCAGAAAACACTT | GUGUUUUCUGGAUAGAGGATT |
| **siSOD1-201** | UCUAUCCAGAAAACACGGUTT | ACCGUGUUUUCUGGAUAGATT |
| **siSOD1-208** | AGAAAACACGGUGGGCCAATT | UUGGCCCACCGUGUUUUCUTT |
| **siSOD1-209** | GAAAACACGGUGGGCCAAATT | UUUGGCCCACCGUGUUUUCTT |
| **siSOD1-210** | AAAACACGGUGGGCCAAAGTT | CUUUGGCCCACCGUGUUUUTT |
| **siSOD1-211** | AAACACGGUGGGCCAAAGGTT | CCUUUGGCCCACCGUGUUUTT |
| **siSOD1-212** | AACACGGUGGGCCAAAGGATT | UCCUUUGGCCCACCGUGUUTT |
| **siSOD1-213** | ACACGGUGGGCCAAAGGAUTT | AUCCUUUGGCCCACCGUGUTT |
| **siSOD1-215** | ACGGUGGGCCAAAGGAUGATT | UCAUCCUUUGGCCCACCGUTT |
| **siSOD1-216** | CGGUGGGCCAAAGGAUGAATT | UUCAUCCUUUGGCCCACCGTT |
| **siSOD1-218** | GUGGGCCAAAGGAUGAAGATT | UCUUCAUCCUUUGGCCCACTT |
| **siSOD1-219** | UGGGCCAAAGGAUGAAGAGTT | CUCUUCAUCCUUUGGCCCATT |
| **siSOD1-220** | GGGCCAAAGGAUGAAGAGATT | UCUCUUCAUCCUUUGGCCCTT |
| **siSOD1-222** | GCCAAAGGAUGAAGAGAGGTT | CCUCUCUUCAUCCUUUGGCTT |
| **siSOD1-224** | CAAAGGAUGAAGAGAGGCATT | UGCCUCUCUUCAUCCUUUGTT |
| **siSOD1-225** | AAAGGAUGAAGAGAGGCAUTT | AUGCCUCUCUUCAUCCUUUTT |
| **siSOD1-227** | AGGAUGAAGAGAGGCAUGUTT | ACAUGCCUCUCUUCAUCCUTT |
| **siSOD1-228** | GGAUGAAGAGAGGCAUGUUTT | AACAUGCCUCUCUUCAUCCTT |
| **siSOD1-229** | GAUGAAGAGAGGCAUGUUGTT | CAACAUGCCUCUCUUCAUCTT |
| **siSOD1-231** | UGAAGAGAGGCAUGUUGGATT | UCCAACAUGCCUCUCUUCATT |
| **siSOD1-233** | AAGAGAGGCAUGUUGGAGATT | UCUCCAACAUGCCUCUCUUTT |
| **siSOD1-234** | AGAGAGGCAUGUUGGAGACTT | GUCUCCAACAUGCCUCUCUTT |
| **siSOD1-235** | GAGAGGCAUGUUGGAGACUTT | AGUCUCCAACAUGCCUCUCTT |
| **siSOD1-236** | AGAGGCAUGUUGGAGACUUTT | AAGUCUCCAACAUGCCUCUTT |
| **siSOD1-237** | GAGGCAUGUUGGAGACUUGTT | CAAGUCUCCAACAUGCCUCTT |
| **siSOD1-241** | CAUGUUGGAGACUUGGGCATT | UGCCCAAGUCUCCAACAUGTT |
| **siSOD1-242** | AUGUUGGAGACUUGGGCAATT | UUGCCCAAGUCUCCAACAUTT |
| **siSOD1-243** | UGUUGGAGACUUGGGCAAUTT | AUUGCCCAAGUCUCCAACATT |

| | | |
|---|---|---|
| **siSOD1-244** | GUUGGAGACUUGGGCAAUGTT | CAUUGCCCAAGUCUCCAACTT |
| **siSOD1-245** | UUGGAGACUUGGGCAAUGUTT | ACAUUGCCCAAGUCUCCAATT |
| **siSOD1-246** | UGGAGACUUGGGCAAUGUGTT | CACAUUGCCCAAGUCUCCATT |
| **siSOD1-247** | GGAGACUUGGGCAAUGUGATT | UCACAUUGCCCAAGUCUCCTT |
| **siSOD1-249** | AGACUUGGGCAAUGUGACUTT | AGUCACAUUGCCCAAGUCUTT |
| **siSOD1-252** | CUUGGGCAAUGUGACUGCUTT | AGCAGUCACAUUGCCCAAGTT |
| **siSOD1-254** | UGGGCAAUGUGACUGCUGATT | UCAGCAGUCACAUUGCCCATT |
| **siSOD1-256** | GGCAAUGUGACUGCUGACATT | UGUCAGCAGUCACAUUGCCTT |
| **siSOD1-257** | GCAAUGUGACUGCUGACAATT | UUGUCAGCAGUCACAUUGCTT |
| **siSOD1-258** | CAAUGUGACUGCUGACAAATT | UUUGUCAGCAGUCACAUUGTT |
| **siSOD1-260** | AUGUGACUGCUGACAAAGATT | UCUUUGUCAGCAGUCACAUTT |
| **siSOD1-261** | UGUGACUGCUGACAAAGAUTT | AUCUUUGUCAGCAGUCACATT |
| **siSOD1-262** | GUGACUGCUGACAAAGAUGTT | CAUCUUUGUCAGCAGUCACTT |
| **siSOD1-263** | UGACUGCUGACAAAGAUGCTT | GCAUCUUUGUCAGCAGUCATT |
| **siSOD1-264** | GACUGCUGACAAAGAUGCUTT | AGCAUCUUUGUCAGCAGUCTT |
| **siSOD1-266** | CUGCUGACAAAGAUGCUGUTT | ACAGCAUCUUUGUCAGCAGTT |
| **siSOD1-270** | UGACAAAGAUGCUGUGGCCTT | GGCCACAGCAUCUUUGUCATT |
| **siSOD1-272** | ACAAAGAUGCUGUGGCCGATT | UCGGCCACAGCAUCUUUGUTT |
| **siSOD1-273** | CAAAGAUGCUGUGGCCGAUTT | AUCGGCCACAGCAUCUUUGTT |
| **siSOD1-275** | AAGAUGCUGUGGCCGAUGUTT | ACAUCGGCCACAGCAUCUUTT |
| **siSOD1-276** | AGAUGCUGUGGCCGAUGUGTT | CACAUCGGCCACAGCAUCUTT |
| **siSOD1-279** | UGCUGUGGCCGAUGUGUCUTT | AGACACAUCGGCCACAGCATT |
| **siSOD1-280** | GCUGUGGCCGAUGUGUCUATT | UAGACACAUCGGCCACAGCTT |
| **siSOD1-281** | CUGUGGCCGAUGUGUCUAUTT | AUAGACACAUCGGCCACAGTT |
| **siSOD1-282** | UGUGGCCGAUGUGUCUAUUTT | AAUAGACACAUCGGCCACATT |
| **siSOD1-283** | GUGGCCGAUGUGUCUAUUGTT | CAAUAGACACAUCGGCCACTT |
| **siSOD1-284** | UGGCCGAUGUGUCUAUUGATT | UCAAUAGACACAUCGGCCATT |
| **siSOD1-285** | GGCCGAUGUGUCUAUUGAATT | UUCAAUAGACACAUCGGCCTT |
| **siSOD1-286** | GCCGAUGUGUCUAUUGAAGTT | CUUCAAUAGACACAUCGGCTT |
| **siSOD1-287** | CCGAUGUGUCUAUUGAAGATT | UCUUCAAUAGACACAUCGGTT |
| **siSOD1-288** | CGAUGUGUCUAUUGAAGAUTT | AUCUUCAAUAGACACAUCGTT |
| **siSOD1-299** | UUGAAGAUUCUGUGAUCUCTT | GAGAUCACAGAAUCUUCAATT |
| **siSOD1-300** | UGAAGAUUCUGUGAUCUCATT | UGAGAUCACAGAAUCUUCATT |
| **siSOD1-301** | GAAGAUUCUGUGAUCUCACTT | GUGAGAUCACAGAAUCUUCTT |
| **siSOD1-302** | AAGAUUCUGUGAUCUCACUTT | AGUGAGAUCACAGAAUCUUTT |
| **siSOD1-303** | AGAUUCUGUGAUCUCACUCTT | GAGUGAGAUCACAGAAUCUTT |
| **siSOD1-304** | GAUUCUGUGAUCUCACUCUTT | AGAGUGAGAUCACAGAAUCTT |
| **siSOD1-305** | AUUCUGUGAUCUCACUCUCTT | GAGAGUGAGAUCACAGAAUTT |
| **siSOD1-306** | UUCUGUGAUCUCACUCUCATT | UGAGAGUGAGAUCACAGAATT |
| **siSOD1-307** | UCUGUGAUCUCACUCUCAGTT | CUGAGAGUGAGAUCACAGATT |
| **siSOD1-309** | UGUGAUCUCACUCUCAGGATT | UCCUGAGAGUGAGAUCACATT |
| **siSOD1-311** | UGAUCUCACUCUCAGGAGATT | UCUCCUGAGAGUGAGAUCATT |
| **siSOD1-314** | UCUCACUCUCAGGAGACCATT | UGGUCUCCUGAGAGUGAGATT |
| **siSOD1-315** | CUCACUCUCAGGAGACCAUTT | AUGGUCUCCUGAGAGUGAGTT |
| **siSOD1-316** | UCACUCUCAGGAGACCAUUTT | AAUGGUCUCCUGAGAGUGATT |
| **siSOD1-317** | CACUCUCAGGAGACCAUUGTT | CAAUGGUCUCCUGAGAGUGTT |
| **siSOD1-318** | ACUCUCAGGAGACCAUUGCTT | GCAAUGGUCUCCUGAGAGUTT |

| | | |
|---|---|---|
| **siSOD1-319** | CUCUCAGGAGACCAUUGCATT | UGCAAUGGUCUCCUGAGAGTT |
| **siSOD1-320** | UCUCAGGAGACCAUUGCAUTT | AUGCAAUGGUCUCCUGAGATT |
| **siSOD1-322** | UCAGGAGACCAUUGCAUCATT | UGAUGCAAUGGUCUCCUGATT |
| **siSOD1-323** | CAGGAGACCAUUGCAUCAUTT | AUGAUGCAAUGGUCUCCUGTT |
| **siSOD1-324** | AGGAGACCAUUGCAUCAUUTT | AAUGAUGCAAUGGUCUCCUTT |
| **siSOD1-325** | GGAGACCAUUGCAUCAUUGTT | CAAUGAUGCAAUGGUCUCCTT |
| **siSOD1-326** | GAGACCAUUGCAUCAUUGGTT | CCAAUGAUGCAAUGGUCUCTT |
| **siSOD1-327** | AGACCAUUGCAUCAUUGGCTT | GCCAAUGAUGCAAUGGUCUTT |
| **siSOD1-333** | UUGCAUCAUUGGCCGCACATT | UGUGCGGCCAAUGAUGCAATT |
| **siSOD1-335** | GCAUCAUUGGCCGCACACUTT | AGUGUGCGGCCAAUGAUGCTT |
| **siSOD1-338** | UCAUUGGCCGCACACUGGUTT | ACCAGUGUGCGGCCAAUGATT |
| **siSOD1-344** | GCCGCACACUGGUGGUCCATT | UGGACCACCAGUGUGCGGCTT |
| **siSOD1-347** | GCACACUGGUGGUCCAUGATT | UCAUGGACCACCAGUGUGCTT |
| **siSOD1-348** | CACACUGGUGGUCCAUGAATT | UUCAUGGACCACCAGUGUGTT |
| **siSOD1-349** | ACACUGGUGGUCCAUGAAATT | UUUCAUGGACCACCAGUGUTT |
| **siSOD1-350** | CACUGGUGGUCCAUGAAAATT | UUUUCAUGGACCACCAGUGTT |
| **siSOD1-351** | ACUGGUGGUCCAUGAAAAATT | UUUUUCAUGGACCACCAGUTT |
| **siSOD1-352** | CUGGUGGUCCAUGAAAAAGTT | CUUUUUCAUGGACCACCAGTT |
| **siSOD1-353** | UGGUGGUCCAUGAAAAAGCTT | GCUUUUUCAUGGACCACCATT |
| **siSOD1-354** | GGUGGUCCAUGAAAAAGCATT | UGCUUUUUCAUGGACCACCTT |
| **siSOD1-356** | UGGUCCAUGAAAAAGCAGATT | UCUGCUUUUUCAUGGACCATT |
| **siSOD1-357** | GGUCCAUGAAAAAGCAGAUTT | AUCUGCUUUUUCAUGGACCTT |
| **siSOD1-359** | UCCAUGAAAAAGCAGAUGATT | UCAUCUGCUUUUUCAUGGATT |
| **siSOD1-360** | CCAUGAAAAAGCAGAUGACTT | GUCAUCUGCUUUUUCAUGGTT |
| **siSOD1-361** | CAUGAAAAAGCAGAUGACUTT | AGUCAUCUGCUUUUUCAUGTT |
| **siSOD1-363** | UGAAAAAGCAGAUGACUUGTT | CAAGUCAUCUGCUUUUUCATT |
| **siSOD1-365** | AAAAAGCAGAUGACUUGGGTT | CCCAAGUCAUCUGCUUUUUTT |
| **siSOD1-366** | AAAAGCAGAUGACUUGGGCTT | GCCCAAGUCAUCUGCUUUUTT |
| **siSOD1-367** | AAAGCAGAUGACUUGGGCATT | UGCCCAAGUCAUCUGCUUUTT |
| **siSOD1-368** | AAGCAGAUGACUUGGGCAATT | UUGCCCAAGUCAUCUGCUUTT |
| **siSOD1-369** | AGCAGAUGACUUGGGCAAATT | UUUGCCCAAGUCAUCUGCUTT |
| **siSOD1-370** | GCAGAUGACUUGGGCAAAGTT | CUUUGCCCAAGUCAUCUGCTT |
| **siSOD1-371** | CAGAUGACUUGGGCAAAGGTT | CCUUUGCCCAAGUCAUCUGTT |
| **siSOD1-372** | AGAUGACUUGGGCAAAGGUTT | ACCUUUGCCCAAGUCAUCUTT |
| **siSOD1-375** | UGACUUGGGCAAAGGUGGATT | UCCACCUUUGCCCAAGUCATT |
| **siSOD1-376** | GACUUGGGCAAAGGUGGAATT | UUCCACCUUUGCCCAAGUCTT |
| **siSOD1-377** | ACUUGGGCAAAGGUGGAAATT | UUUCCACCUUUGCCCAAGUTT |
| **siSOD1-378** | CUUGGGCAAAGGUGGAAAUTT | AUUUCCACCUUUGCCCAAGTT |
| **siSOD1-379** | UUGGGCAAAGGUGGAAAUGTT | CAUUUCCACCUUUGCCCAATT |
| **siSOD1-380** | UGGGCAAAGGUGGAAAUGATT | UCAUUUCCACCUUUGCCCATT |
| **siSOD1-381** | GGGCAAAGGUGGAAAUGAATT | UUCAUUUCCACCUUUGCCCTT |
| **siSOD1-382** | GGCAAAGGUGGAAAUGAAGTT | CUUCAUUUCCACCUUUGCCTT |
| **siSOD1-383** | GCAAAGGUGGAAAUGAAGATT | UCUUCAUUUCCACCUUUGCTT |
| **siSOD1-384** | CAAAGGUGGAAAUGAAGAATT | UUCUUCAUUUCCACCUUUGTT |
| **siSOD1-386** | AAGGUGGAAAUGAAGAAAGTT | CUUUCUUCAUUUCCACCUUTT |
| **siSOD1-387** | AGGUGGAAAUGAAGAAAGUTT | ACUUUCUUCAUUUCCACCUTT |
| **siSOD1-388** | GGUGGAAAUGAAGAAAGUATT | UACUUUCUUCAUUUCCACCTT |

| | | |
|---|---|---|
| **siSOD1-389** | GUGGAAAUGAAGAAAGUACTT | GUACUUUCUUCAUUUCCACTT |
| **siSOD1-397** | GAAGAAAGUACAAAGACAGTT | CUGUCUUUGUACUUUCUUCTT |
| **siSOD1-399** | AGAAAGUACAAAGACAGGATT | UCCUGUCUUUGUACUUUCUTT |
| **siSOD1-400** | GAAAGUACAAAGACAGGAATT | UUCCUGUCUUUGUACUUUCTT |
| **siSOD1-402** | AAGUACAAAGACAGGAAACTT | GUUUCCUGUCUUUGUACUUTT |
| **siSOD1-405** | UACAAAGACAGGAAACGCUTT | AGCGUUUCCUGUCUUUGUATT |
| **siSOD1-408** | AAAGACAGGAAACGCUGGATT | UCCAGCGUUUCCUGUCUUUTT |
| **siSOD1-409** | AAGACAGGAAACGCUGGAATT | UUCCAGCGUUUCCUGUCUUTT |
| **siSOD1-410** | AGACAGGAAACGCUGGAAGTT | CUUCCAGCGUUUCCUGUCUTT |
| **siSOD1-411** | GACAGGAAACGCUGGAAGUTT | ACUUCCAGCGUUUCCUGUCTT |
| **siSOD1-412** | ACAGGAAACGCUGGAAGUCTT | GACUUCCAGCGUUUCCUGUTT |
| **siSOD1-414** | AGGAAACGCUGGAAGUCGUTT | ACGACUUCCAGCGUUUCCUTT |
| **siSOD1-415** | GGAAACGCUGGAAGUCGUUTT | AACGACUUCCAGCGUUUCCTT |
| **siSOD1-416** | GAAACGCUGGAAGUCGUUUTT | AAACGACUUCCAGCGUUUCTT |
| **siSOD1-417** | AAACGCUGGAAGUCGUUUGTT | CAAACGACUUCCAGCGUUUTT |
| **siSOD1-418** | AACGCUGGAAGUCGUUUGGTT | CCAAACGACUUCCAGCGUUTT |
| **siSOD1-420** | CGCUGGAAGUCGUUUGGCUTT | AGCCAAACGACUUCCAGCGTT |
| **siSOD1-421** | GCUGGAAGUCGUUUGGCUUTT | AAGCCAAACGACUUCCAGCTT |
| **siSOD1-423** | UGGAAGUCGUUUGGCUUGUTT | ACAAGCCAAACGACUUCCATT |
| **siSOD1-424** | GGAAGUCGUUUGGCUUGUGTT | CACAAGCCAAACGACUUCCTT |
| **siSOD1-425** | GAAGUCGUUUGGCUUGUGGTT | CCACAAGCCAAACGACUUCTT |
| **siSOD1-426** | AAGUCGUUUGGCUUGUGGUTT | ACCACAAGCCAAACGACUUTT |
| **siSOD1-429** | UCGUUUGGCUUGUGGUGUATT | UACACCACAAGCCAAACGATT |
| **siSOD1-430** | CGUUUGGCUUGUGGUGUAATT | UUACACCACAAGCCAAACGTT |
| **siSOD1-431** | GUUUGGCUUGUGGUGUAAUTT | AUUACACCACAAGCCAAACTT |
| **siSOD1-432** | UUUGGCUUGUGGUGUAAUUTT | AAUUACACCACAAGCCAAATT |
| **siSOD1-433** | UUGGCUUGUGGUGUAAUUGTT | CAAUUACACCACAAGCCAATT |
| **siSOD1-434** | UGGCUUGUGGUGUAAUUGGTT | CCAAUUACACCACAAGCCATT |
| **siSOD1-435** | GGCUUGUGGUGUAAUUGGGTT | CCCAAUUACACCACAAGCCTT |
| **siSOD1-436** | GCUUGUGGUGUAAUUGGGATT | UCCCAAUUACACCACAAGCTT |
| **siSOD1-437** | CUUGUGGUGUAAUUGGGAUTT | AUCCCAAUUACACCACAAGTT |
| **siSOD1-443** | GUGUAAUUGGGAUCGCCCATT | UGGGCGAUCCCAAUUACACTT |
| **siSOD1-444** | UGUAAUUGGGAUCGCCCAATT | UUGGGCGAUCCCAAUUACATT |
| **siSOD1-445** | GUAAUUGGGAUCGCCCAAUTT | AUUGGGCGAUCCCAAUUACTT |
| **siSOD1-446** | UAAUUGGGAUCGCCCAAUATT | UAUUGGGCGAUCCCAAUUATT |
| **siSOD1-447** | AAUUGGGAUCGCCCAAUAATT | UUAUUGGGCGAUCCCAAUUTT |

*NOTE:* TT, dual deoxythymidine overhangs

**Supplementary Table 4.** Chemically modified siRNAs, controls, and primer sequences.

| siRNA | Sense (5'-3') | Antisense (5'-3') |
|---|---|---|
| siCON/Neg Con | ACUACUGAGUGACAGUAGAtt | UCUACUGUCACUCAGUAGUtt |
| siSOD1-063M1 | mG*fA*mGfCmAfGmAfAfGfGfAfAmAfGmUfA*mA*fU | mA*fU*mUfAmCfUmUfUmCfCmUfUmCfUmGfCmUfC*mG*fA |
| siSOD1-063M2 | fC*mG*fAmGfCmAfGmAfAfGfGmAfAmAfGmUfA*mA*fU | mA*fU*mUfAmCfUmUfUmCfCmUfUmCfUmGfCmUfCmG*fA*mA |
| siSOD1-063M3 | mU*fC*mGfAmGfCmAfGfAfAfGfGmAfAmAfGmUfA*mA*fU | mA*fU*mUfAmCfUmUfUmCfCmUfUmCfUmGfCmUfCmGfA*mA*fA |
| siSOD1-063M4 | fU*mU*fCmGfAmGfCmAfGfAfAmGfGmAfAmAfGmUfA*mA*fU | mA*fU*mUfAmCfUmUfUmCfCmUfUmCfUmGfCmUfCmGfAmA*fA*mU |
| siSOD1-047M1 | mG*fG*mGfCmAfUmCfAfUfCfAfAmUfUmUfC*mG*fA | mU*fC*mGfAmAfAmUfUmGfAmUfGmAfUmGfCmCfC*mU*fG |
| siSOD1-047M2 | fA*mG*fGmGfCmAfUmCfAfUfCmAfAmUfUmUfC*mG*fA | mU*fC*mGfAmAfAmUfUmGfAmUfGmAfUmGfCmCfCmU*fG*mC |
| siSOD1-047M3 | mC*fA*mGfGmGfCmAfUfCfAfUfCmAfAmUfUmUfC*mG*fA | mU*fC*mGfAmAfAmUfUmGfAmUfGmAfUmGfCmCfCmUfG*mC*fA |
| siSOD1-047M4 | fG*mC*fAmGfGmGfCmAfUfCfAmUfCmAfAmUfUmUfC*mG*fA | mU*fC*mGfAmAfAmUfUmGfAmUfGmAfUmGfCmCfCmUfGmC*fA*mC |
| siSOD1-104M1 | mC*fA*mUfUmAfAmAfGfGfAfCfUmGfAmCfU*mG*fA | mU*fC*mAfGmUfCmAfGmUfCmCfUmUfUmAfAmUfG*mC*fU |
| siSOD1-104M2 | fG*mC*fAmUfUmAfAmAfGfGfAmCfUmGfAmCfU*mG*fA | mU*fC*mAfGmUfCmAfGmUfCmCfUmUfUmAfAmUfGmC*fU*mU |
| siSOD1-104M3 | mA*fG*mCfAmUfUmAfAfAfGfGfAmCfUmGfAmCfU*mG*fA | mU*fC*mAfGmUfCmAfGmUfCmCfUmUfUmAfAmUfGmCfU*mU*fC |
| siSOD1-104M4 | fA*mA*fGmCfAmUfUmAfAfAfGmGfAmCfUmGfAmCfU*mG*fA | mU*fC*mAfGmUfCmAfGmUfCmCfUmUfUmAfAmUfGmCfUmU*fC*mC |
| siSOD1-005M1 | mG*fA*mCfGmAfAmGfGfCfCfGfUmGfUmGfC*mG*fU | mA*fC*mGfCmAfCmAfCmGfGmCfCmUfUmCfGmUfC*mG*fC |
| siSOD1-005M2 | fC*mG*fAmCfGmAfAmGfGfCfCmGfUmGfUmGfC*mG*fU | mA*fC*mGfCmAfCmAfCmGfGmCfCmUfUmCfGmUfCmG*fC*mC |
| siSOD1-005M3 | mG*fC*mGfAmCfGmAfAfGfGfCfCmGfUmGfUmGfC*mG*fU | mA*fC*mGfCmAfCmAfCmGfGmCfCmUfUmCfGmUfCmGfC*mC*fA |
| siSOD1-005M4 | fG*mG*fCmGfAmCfGmAfAfGfGmCfCmGfUmGfUmGfC*mG*fU | mA*fC*mGfCmAfCmAfCmGfGmCfCmUfUmCfGmUfCmGfCmC*fA*mU |
| siSOD1-258M1 | mA*fA*mUfGmUfGmAfCfUfGfCfUmGfAmCfA*mA*fA | mU*fU*mUfGmUfCmAfGmCfAmGfUmCfAmCfAmUfU*mG*fC |
| siSOD1-258M2 | fC*mA*fAmUfGmUfGmAfCfUfGmCfUmGfAmCfA*mA*fA | mU*fU*mUfGmUfCmAfGmCfAmGfUmCfAmCfAmUfUmG*fC*mC |
| siSOD1-258M3 | mG*fC*mAfAmUfGmUfGfAfCfUfGmCfUmGfAmCfA*mA*fA | mU*fU*mUfGmUfCmAfGmCfAmGfUmCfAmCfAmUfUmGfC*mC*fC |
| siSOD1-258M4 | fG*mG*fCmAfAmUfGmUfGfAfCmUfGmCfUmGfAmCfA*mA*fA | mU*fU*mUfGmUfCmAfGmCfAmGfUmCfAmCfAmUfUmGfCmC*fC*mA |
| siSOD1-047M3-AC1 | mC*fA*mGfGmGfCmAfUfCfAfUfCmAfAmUfUmUfC*mG*fA-L9-meU*meU*meG*meU*meA*meU*meU*meC*meU*meA*meU*meG*meU*meU | mU*fC*mGfAmAfAmUfUmGfAmUfGmAfUmGfCmCfCmUfG*mC*fA |
| siSOD1-005M3-AC1 | mG*fC*mGfAmCfGmAfAfGfGfCfCmGfUmGfUmGfC*mG*fU-L9-meU*meU*meG*meU*meA*meU*meU*meC*meU*meA*meU*meG*meU*meU | mA*fC*mGfCmAfCmAfCmGfGmCfCmUfUmCfGmUfCmGfC*mC*fA |
| siCON1-AC1$^{VP}$ | fC*mG*fGmUfGmGfAmGfGfCfCfUfGmGfCmGfCmA*fA*mU-L9-meU*meU*meG*meU*meA*meU*meU*meC*meU*meA*meU*meG*meU*meU | VpmG*fC*mCfAmCfCmUfCmCfGmGfAmCfCmGfCmGfUmUfA*mC*fA |
| siCON2-AC1$^{VP}$ | mA*fG*mUfUmCfGmUfGfUfCfAfUmCfCmAfGmAfG*mA*fU-L9-meU*meU*meG*meU*meA*meU*meU*meC*meU*meA*meU*meG*meU*meU | VpmA*fU*mCfUmCfUmGfGmAfUmGfAmCfAmCfGmAfAmCfU*mU*fG |
| siCON3-AC1$^{VP}$ | mG*fU*mAfCmUfUmUfUfGfUfGfUmAfGmUfAmCfA*mA*fA-L9-meU*meU*meG*meU*meA*meU*meU*meC*meU*meA*meU*meG*meU*meU | VpmU*fU*mUfGmUfAmCfUmAfCmAfCmAfAmAfAmGfUmAfC*mU*fG |
| siSOD1-063M3-AC1$^{VP}$ | mU*fC*mGfAmGfCmAfGfAfAfGfGmAfAmAfGmUfA*mA*fU-L9-meU*meU*meG*meU*meA*meU*meU*meC*meU*meA*meU*meG*meU*meU | VpmA*fU*mUfAmCfUmUfUmCfCmUfUmCfUmGfCmUfCmGfA*mA*fA |
| siSOD1-047M3-AC1$^{VP}$ | mC*fA*mGfGmGfCmAfUfCfAfUfCmAfAmUfUmUfC*mG*fA-L9-meU*meU*meG*meU*meA*meU*meU*meC*meU*meA*meU*meG*meU*meU | VpmU*fC*mGfAmAfAmUfUmGfAmUfGmAfUmGfCmCfCmUfG*mC*fA |
| siSOD1-104M3-AC1$^{VP}$ | mA*fG*mCfAmUfUmAfAfAfGfGfAmCfUmGfAmCfU*mG*fA-L9-meU*meU*meG*meU*meA*meU*meU*meC*meU*meA*meU*meG*meU*meU | VpmU*fC*mAfGmUfCmAfGmUfCmCfUmUfUmAfAmUfGmCfU*mU*fC |
| siSOD1-005M3-AC1$^{VP}$ | mG*fC*mGfAmCfGmAfAfGfGfCfCmGfUmGfUmGfC*mG*fU-L9-meU*meU*meG*meU*meA*meU*meU*meC*meU*meA*meU*meG*meU*meU | VpmA*fC*mGfCmAfCmAfCmGfGmCfCmUfUmCfGmUfCmGfC*mC*fA |
| siSOD1-258M3-AC1$^{VP}$ | mG*fC*mAfAmUfGmUfGfAfCfUfGmCfUmGfAmCfA*mA*fA-L9-meU*meU*meG*meU*meA*meU*meU*meC*meU*meA*meU*meG*meU*meU | VpmU*fU*mUfGmUfCmAfGmCfAmGfUmCfAmCfAmUfUmGfC*mC*fC |
| siSOD1-047M3$^{VP}$ | mC*fA*mGfGmGfCmAfUfCfAfUfCmAfAmUfUmUfC*mG*fA | VpmU*fC*mGfAmAfAmUfUmGfAmUfGmAfUmGfCmCfCmUfG*mC*fA |
| siSOD1-005M3$^{VP}$ | mG*fC*mGfAmCfGmAfAfGfGfCfCmGfUmGfUmGfC*mG*fU | VpmA*fC*mGfCmAfCmAfCmGfGmCfCmUfUmCfGmUfCmGfC*mC*fA |
| **ASO/ACO** | **Sequence (5'-3')** | |
| ASO$^{SOD1}$ | meC*meAmeG*meGmeA*t*a*c*a*t*t*t*c*t*a*meCmeA*meGmeC*meT | |
| AC1 | L9-meU*meU*meG*meU*meA*meU*meU*meC*meU*meA*meU*meG*meU*meU | |
| **Primers** | **Forward (5'-3')** | **Reverse (5'-3')** |
| SOD1 | aagcattaaaggactgactgaagg | caagtctccaacatgcctctc |
| TBP (human) | tgctcacccaccaacaatttag | tctgctctgactttagcacctg |
| Tbp (mouse) | ccgtgaatcttggctgtaaact | tgtccgtggctctcttattc |

f: 2'-fluoro, m: 2'-O-methyl, *: phosphorothioate, Vp: 5'-(*E*)-vinylphosphonate, L9: PEG linker, me: 2'-O-methoxy-ethyl, meC: 2'-O-methoxyethyl-5-methylcytosine, meU: 2'-O-methoxyethyl-5-methyl-uridine, c: 5-methyl-2'-deoxycytidine, lower case: DNA
